# Evolutionary epidemiology consequences of trait-dependent control of heterogeneous parasites

**DOI:** 10.1101/2021.06.08.447562

**Authors:** Leonardo Miele, R M L Evans, Nik Cunniffe, Clara Torres-Barcelo, Daniele Bevacqua

## Abstract

Disease control can induce both demographic and evolutionary responses in host-parasite systems. Foreseeing the outcome of control therefore requires knowledge of the eco-evolutionary feedback between control and system. Previous work has assumed that control strategies have a homogeneous effect on the parasite population. However, this is not true when control targets those traits that confer to the parasite heterogeneous levels of resistance, which can additionally be related to other key parasite traits, through evolutionary trade-offs. In this work, we develop a minimal model coupling epidemiological and evolutionary dynamics to explore possible trait-dependent effects of control strategies. Particularly, we consider a parasite expressing continuous levels of a trait determining resource exploitation, and a control treatment that can be either positively or negatively correlated with that trait. We demonstrate the potential of trait-dependent control by considering that the decision maker may want to minimise both the damage caused by the disease and the use of treatment, due to possible environmental or economic costs. We identify efficient strategies showing that the optimal type of treatment depends on the amount applied. Our results pave the way for the study of control strategies based on evolutionary constraints, such as collateral sensitivity and resistance costs, which are receiving increasing attention for both public health and agricultural purposes.

## Introduction

Disease and pest control strategies aim to eradicate or mitigate the exploitation of a parasite population, on a host population of economic (agriculture) or public health (humans) interest (Gilligan, 2002). By altering the ecological host-parasite interactions, a control strategy can induce both demographic effects (Gilligan and van den Bosch, 2008) and evolutionary responses (Day et al., 2020). Predicting the outcome of control strategies on a system therefore depends on our understanding of its eco-evolutionary feedbacks (Day and Gandon, 2006), that is on the evolutionary epidemiology behaviour (Galvani, 2003).

In addition to experimental studies, theoretical work based on evolutionary epidemiology has provided insights into control strategies and on their, often counter-intuitive, consequences. For instance, host reduction, *e*.*g*. via culling, may increase disease abundance and prevalence, (Bolzoni and De Leo, 2013); altering parasite growth, such as in vaccination campaigns, can lead to selection for more virulent parasites, thus increasing disease severity (Zurita-Gutiérrez and Lion, 2015; Gandon et al., 2003, 2001). Studies on the emergence and spread of multidrug resistance have also shown that the efficacy of a control strategy may depend on the structure of the host population (*e*.*g*. age or spatial distributions), as well as on parasite heterogeneity (McLeod and Gandon, 2021; Lehtinen et al., 2019; Blanquart et al., 2018). Furthermore, models also provide a low-cost tool to explore and optimise large-scale agricultural practices (Rimbaud et al., 2018), such as the deployment of disease-resistant cultivars (Taylor and Cunniffe, 2022) and crop rotation (Bargués-Ribera and Gokhale, 2020), whose in-field implementation can consume considerable time and resources.

Most of the experimental and theoretical work has assumed that control strategies have an homogeneous effect on parasites, or that parasites are endowed with a genetically encoded resistance that is either present or absent (qualitative resistance) (consortium REX, 2010). However, this picture neglects the cases where the efficacy of a treatment depends on quantitative traits that can be heterogeneously expressed in the parasite population (quantitative resistance) (Corwin and Kliebenstein, 2017). In fact, parasites have developed several defence mechanisms that can quantitatively affect drug uptake, leading to variable levels of resistance (El Meouche and Dunlop, 2018; Munita and Arias, 2016). Widespread examples of such mechanisms are: metabolic regulators (Chebotar et al., 2021), biofilms (Fanning and Mitchell, 2012; Costerton et al., 1999), efflux pumps (Martinez et al., 2009), flagella (Lyu et al., 2021) and capsules (Song et al., 2021). Thus, a parasite population will often exhibit heterogeneity in the expression of key traits (Dutta et al., 2020; González et al., 2019; Perrier et al., 2019; Schrö ter and Dersch, 2019; Hewitt et al., 2016), potentially leading to heterogeneous trait-dependent treatment effects (Alizon, 2020; Martίnez et al., 2019; Laine and Barrès, 2013; Porco et al., 2005), and ultimately threatening control strategy’s overall efficacy (Dewachter et al., 2019; Weigel and Dersch, 2018; Patyka et al., 2016; Gefen and Balaban, 2009).

The occurrence of trait-dependent treatment effects is expected to be exacerbated in the future, as many of the new promising strategies to counter resistance escalation are based on the exploitation of trait-specific evolutionary constraints (Furusawa et al., 2018; Lässig et al., 2017; Palmer and Kishony, 2013), such as: fitness costs of resistance (Hawkins and Fraaije, 2018; Vincent et al., 2013; Andersson and Hughes, 2010; Lenski, 1998), life-history (Shoval et al., 2012) and metabolic (Pinheiro et al., 2021; Weiße et al., 2015) trade-offs and collateral sensitivity (Roemhild and Andersson, 2021; Maeda et al., 2020; Barbosa et al., 2019; Maltas and Wood, 2019; Lázár et al., 2018). For instance, bacterial efflux pumps rely on proton motive force to both import some toxic compounds (*e*.*g*. aminoglycosides) (Alekshun and Levy, 2007; Taber et al., 1987), and to expel others (*e*.*g*. *β*-lactams) (Suzuki et al., 2014; Lázár et al., 2013; Okusu et al., 1996); therefore, strains with a reduced proton motive force will be more resistant to one antibiotic, but more sensitive to the other, and vice-versa (Roemhild and Andersson, 2021; Pál et al., 2015). This phenomenon is a textbook example of antibiotic collateral sensitivity, but it also affects other types of control strategies, such as fungicides, copper use or phage therapy, and it can potentially involve other fields of application, such as biocontrol methods in agriculture.

Phage therapy employs viruses (phages) that selectively attack bacteria and ultimately kill them. It has been observed that resistance to both phages and antibiotics is often costly for bacteria (Laure and Ahn, 2022; Mangalea and Duerkop, 2020; Meaden et al., 2015): selection pressure often leads to the evolution of bacterial strains that are resistant to either phage or antibiotic therapy, which can therefore be applied in combination to obtain synergistic antimicrobial effects (Coyne et al., 2022; Kebriaei et al., 2022; Torres-Barceló and Hochberg, 2016). In addition, recently discovered phages targeting mechanisms involved in both antibiotic resistance and virulence are smart tools to restore antibiotic treatment efficacy, and co-select for avirulent strains (Gurney et al., 2020; Chan et al., 2018). Phages attach to specific bacterial external structures (*e*.*g*. flagella, capsules, or efflux pumps) that are involved in important biological processes, such as antibiotic resistance and pathogenicity (Esteves and Scharf, 2022; Song et al., 2021; Chan et al., 2016). Modifications in these components make phage infection more difficult and are therefore selected for in bacterial populations exposed to phages. These changes inhibit the bacteria’s previous ability to cause disease and to resist antibiotics, thereby restoring their sensitivity to the treatment (Gurney et al., 2020; Chiarelli et al., 2020; Chan et al., 2016). Specifically, bacteria with a down-regulated production of efflux pumps would avoid phage infection on the one hand. On the other hand, however, they would be more sensitive to antibiotics or toxic heavy metals, that are detoxified by these pumps.

A similar mechanism underlies the behaviour observed in *Myzus persicae*, an aphid considered a major threat to agriculture (Van Emden and Harrington, 2017) and an important living model for the study of insecticide resistance (Bass et al., 2014): mutations in the metabolic activity can lead to the emergence of clones that are more resistant to insecticides due to reduced uptake rates, but also more vulnerable to natural enemies (Foster et al., 2007), with non-trivial consequences on their demography (Foster et al., 2011). When exposed to both insecticides and biocontrol (natural enemies or pathogens), insects may face fitness trade-offs that prevent them from maintaining the same level of resistance (Lacey et al., 2015). Therefore, by exerting different selection pressures on a pest, a synergistic use of chemical and biocontrol has the potential to contain resistance development and maintain crop productivity, while minimising the negative environmental impacts by potentially reducing chemical doses (Ons et al., 2020), akin to the phage-antibiotics case.

Since traits that lead to resistance are often also involved in the parasite’s ability to exploit and harm the host (Copin et al., 2019; Giraud et al., 2017; Alcalde-Rico et al., 2016; Beceiro et al., 2013), the outcome of a trait-specific treatment may be complicated by the presence of trade-offs between resistance mechanisms and other life-history traits of the parasite (Boots and Bowers, 2004; Boots and Haraguchi, 1999), such as those involving host exploitation and disease-induced mortality (known as Transmission-Virulence trade-off) in the case of micro-parasites (Alizon et al., 2009; van den Bosch and Gilligan, 2008; Bull, 1994), and those involving foraging activity and lifespan in the case of macro-parasites (Strobbe et al., 2011; Stoks et al., 2005; Brodin and Johansson, 2004; Gotthard, 2000; Anholt et al., 2000; Werner and Anholt, 1993). Knowledge of novel trait-specific mechanisms is broadening the spectrum of possible selection pressures we can exert on parasites (Allen et al., 2014), and a full exploitation of the potential of such new strategies depends on our understanding of the eco-evolutionary feedbacks between the treatment and the biological system, at various levels of description (Perry, 2021; Burmeister et al., 2021).

Accounting for a detailed description of the therapy-parasite interactions have provided insightful information on the *in vitro* behaviour of specific systems (Aulin et al., 2021; Rodriguez-Gonzalez et al., 2020; Nichol et al., 2019; Mattei et al., 2018; Bull et al., 2014). However, their results can be hardly generalisable, and the corresponding population-level information can be tricky to obtain. Here, we are interested in developing a general framework, shared in principle by any control strategy where: *i*) treatment efficacy depends on a target trait; *ii*) different treatment types correlate differently with the target trait; *iii*) target traits are heterogeneously expressed across the parasite population, leading to heterogeneous treatment effects; *iv*) target traits may be related to other parasite’s traits through trade-offs.

We tackle the above issue by means of a minimal ecological model describing the dynamics of a generic, valuable resource, and of a generic parasite population (Lafferty et al., 2015). Following a well established evolutionary epidemiological approach (Day and Proulx, 2004), we model parasites as a trait-structured population, characterised by heterogeneous levels of expression of key traits. We consider a single proxy trait variable accounting for the possible trade-offs between the parasite’s level of exploitation and mortality, and its resistance to treatment. Crucially and differently from previous work (Alizon and van Baalen, 2005; Porco et al., 2005; Frank, 1996), we allow treatment efficacy to either correlate or anti-correlate with the proxy trait, in order to reproduce potentially different control strategies, as well as combinations of them. We focus our analysis on the implications of trait-dependent treatment from an agricultural perspective, where maximising resources and reducing treatment use are two major objectives (Medina-Pastor and

Triacchini, 2020; Organization et al., 2014). Using a simple multi-criteria analysis, we show that the ability tune trait-dependent control can harness parasites heterogeneity to our advantage: particularly, we show the existence of optimal treatment types, and the emergence of saturation effects on resource production gains. By focusing on a quantity of general, economic interest (healthy resource at equilibrium), our results can be applied to a variety of agricultural practices, and are potentially extendable to different scenarios.

## Models and methods

### The model

In the spirit of Lafferty et al. (2015), we start with a minimal ecological model, describing the interaction between (*R*)esource and (*P*)arasite biomass, potentially representative of both micro-parasite and macro-parasite scenarios. The former scenario neglects the explicit dynamics of within-host abundance (as is commonly done for pathogens) (Keeling and Rohani, 2011; Anderson and May, 1979): *R* represents the biomass of healthy, susceptible hosts, while *P* represents the biomass of hosts infected by the pathogen. In the latter scenario (typical of crop-pest and plant-herbivore systems), *R* represents the resource biomass, and *P* the biomass of the parasite that consumes it. In both scenarios, *R* is treated as a renewable resource exploited by *P*. The treatment has the effect of killing the parasite, thus removing *P* biomass. In what follows, we first present the underlying assumptions of the minimal model, from which we then develop the eco-evolutionary formulation.

### Homogeneous *RP* formulation with treatment

Our assumptions are summarised as follows: *(i)* resource biomass is renewed at a constant rate, and (*ii*) is converted to parasite biomass upon exploitation: (*ii_micro_*) in the micro- scenario, it corresponds to the transmission of the infection from an infected to a susceptible host; (*ii_macro_*) in the macro- scenario, to the consumption of the resource by the consumer. (*iii*) Resource and parasite biomass can be removed from the system, by various possible mechanisms acting on both (*e*.*g*. natural mortality) or either (*e*.*g*. diseaseinduced mortality). (*iv*) Parasite biomass is eradicated at a rate proportional to treatment application and efficacy: (*iv_micro_*) in the micro- scenario, eradication is intended in the sense of removing the pathogen from the host (Castle and Gilligan, 2012; Hall et al., 2004); (*iv_macro_*) in the macroscenario, eradication is intended in the sense of killing the parasite (Van Den Bosch and Gilligan, 2008).

Under the above assumptions, the dynamics of *R* and *P* biomass is given by the following system of equations:

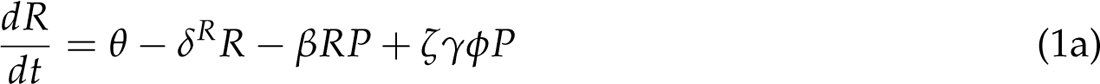

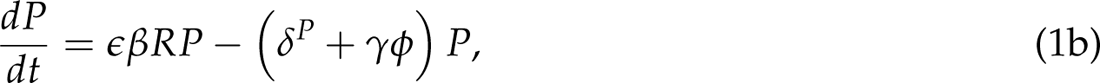

where *θ* is the resource renewal rate; *δ^R^*^,*P*^ are the mortality rates; *β* is the exploitation rate; *ɛ* is the coefficient accounting for biomass conversion; *γϕ* is the total eradication rate per unit of parasite biomass, where parameter *γ* is the treatment application rate, while *ϕ* is the treatment efficacy; *ζ* is the treated parasite’s fate parameter.

In the microscenario, *β* corresponds to the transmission rate of the infection, and *ɛ* = 1; the fate of the treated infected host is determined by *ζ* as follows: for *ζ* = 1, the treated host is restored to *R* (*i*.e. treatment provides recovery without immunity, as it may be the case with animal antibiotics) (Forster and Gilligan, 2007; Hethcote, 2000); for *ζ* = 0, the treated host is removed from the system (recovery with immunity or permanent removal, as is generally the case with plant diseases) (Hall et al., 2004). Intermediate values 0 < ι < ι 1 can model in-between cases. As it will be shown, for our purposes it is not necessary to specify the fate of the treated host, since the results presented here are independent of the value of *ζ*.

In the macro- scenario, *β* corresponds to the consumption rate of the resource, 0 *< ζ <* 1 (biomass conversion from resource to consumer) and *ζ* = 0 (treatment removes consumers from the system).

In the standard ecological formulation of the model, the population is considered to be homogeneous, so that treatment efficacy, exploitation and mortality rates have constant values across individuals. The behaviour of this basic model is well known, and we refer the reader to Korobeinikov and Wake (2002) for details.

### Trait-structured *RPx* formulation with trait-dependent treatment

Following previous work (Sasaki et al., 2021; Day et al., 2020; Korobeinikov, 2018; Day and Proulx, 2004), we elaborate the formulation equivalent to the above model Eqs. (1), in the case of a parasite heterogeneously expressing a trait. The parameters of the model and their description are summarised in Table 1.

**Table 1:**
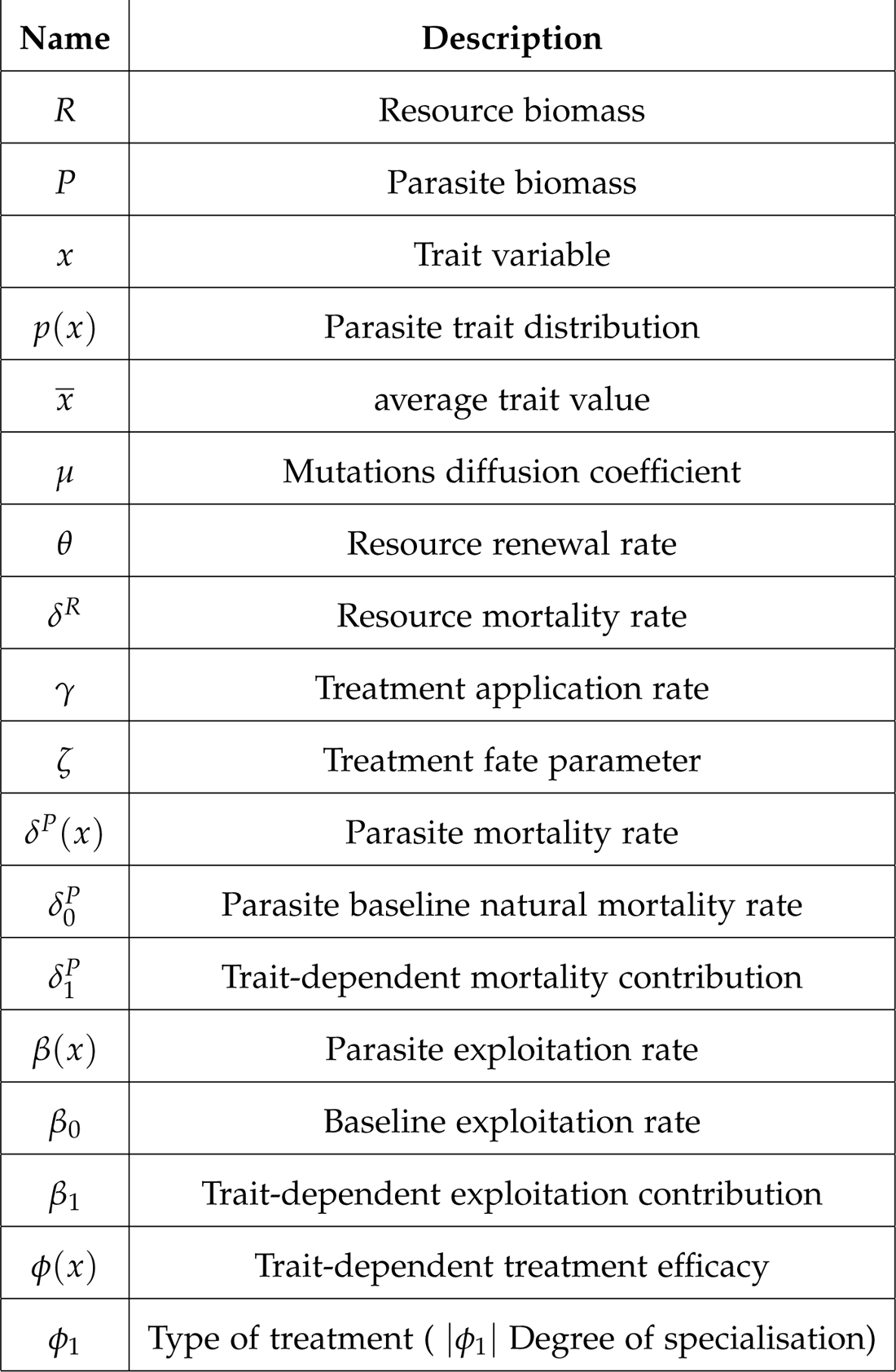
List of the variables and parameters used in the model.

The following assumptions are introduced: (*i*) The parasite has a *continuum-of-strains*, mathematically described by a single continuous proxy variable *x ∈ T*, where *T* is the trait space. For the sake of simplicity, we will also consider the unit interval *T* = [0, 1] as trait space. The generalisation to any positive interval [*x*_min_, *x*_max_] is straightforward, and can be mapped to the interval [0, 1] by rescaling of the parameters. Heterogeneity is described by the trait distribution *p*(*x*), that is the density of parasites carrying trait *x*. (*ii*) The trait determines the parasite’s levels of resource exploitation, mortality and treatment efficacy, which are now represented by the functions *β*(*x*), *δ^P^*(*x*), *ϕ*(*x*). (*iii*) Parasites undergo mutations that induce small changes in their traits, and maintain heterogeneity within the population; mutation rates are high compared to the ecological timescale, they are unbiased and no preferred direction is assumed.

In the heterogeneous formulation, the system is described by the ecological and the evolutionary states. The ecological state is given by the amount of *R* and *P* biomass. The evolutionary state is given by the trait distribution *p*(*x*). Mathematically speaking, *p*(*x*) is a probability distribution, over which is possible to compute average quantities with respect to the parasite population. Given the above assumptions, the dynamics of the heterogeneous system is provided by the following system of equations (details in the Supplemental Material):

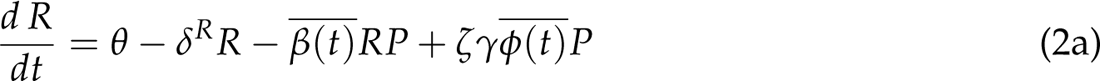

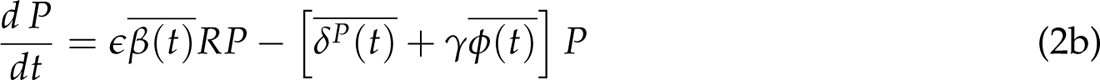

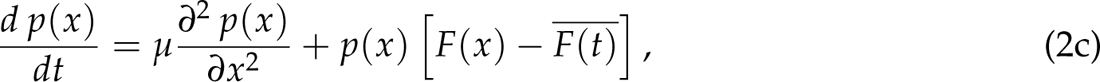

with

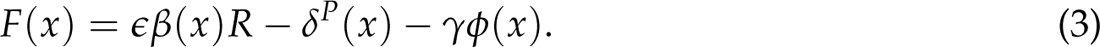

The bar notation indicates the average over the trait distribution, thus 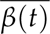 is the average exploitation rate of the population:

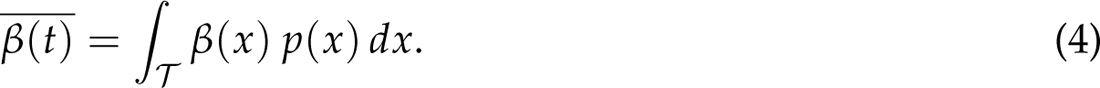

In the above equation, the time dependence is retained in order to recognise that such averages are not fixed, but change over time with the time variation of the trait distribution *p*(*x*). Equivalent definitions apply to average mortality 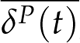, the average treatment efficacy 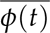 and the function 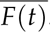.

Eqs. (2a) and (2b) describe the population dynamics at the demographic level. They are equivalent to the classical formulation Eqs. (1), upon replacing the single-strain parameters *β*, *δ^P^*, *ϕ* with their population average counterparts 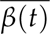, 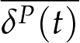, 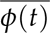. Instead, Eq. (2c) describes the population dynamics at the evolutionary level, as it governs the changes in the parasite trait distribution, due to mutations and competition for limited resource.

Phenotypic mutations are captured by the diffusion operator 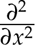 over the trait space, *µ* being the diffusion coefficient related to mutations; this choice assumes that mutations induce small perturbations in the quantitative trait, *i*.*e*. a parasite mutates into a ‘phenotypically close’ variant (possible biases in the direction of mutations may be accounted for, by introducing a gradient term in the last equation) (Lorenzi et al., 2016; Chisholm et al., 2016; Kimura, 1965).

Concomitantly, parasites compete between each other for access to a limited amount of resource (*i*.*e*. infection of a limited number of hosts or consumption of a limited resource), according to the trait-dependent function *F*(*x*): the exploitation term *ɛβ*(*x*)*R* contributes to increasing the density of trait *x*, whereas mortality *δ^P^*(*x*) and efficacy *ϕ*(*x*) contribute to decreasing it.

The overall success of parasites with trait *x* depends on the difference between its value of *F*(*x*) and the population average *F*, as in a replicator dynamics (Schuster and Sigmund, 1983). Thus, *F*(*x*) represents the fitness landscape structuring the parasite’s competition for exploitation. Unlike purely theoretical work, this fitness landscape is not assumed. Rather, it emerges from the ecological interactions (Day et al., 2020). Note that the evolutionary equation takes the same form, regardless of the parasite’s nature and of the fate parameter *ζ*.

The system Eqs. (2) shows neatly the intertwining of ecological and evolutionary levels of description, that is typical of eco-evolutionary dynamics: on the one hand, the demography of the population (given by *R* and *P*) depends on the trait distribution *p*(*x*) via the average quantities 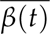, 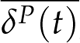, 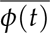; on the other hand, the trait distribution depends on the demography via the ecological interactions (as exploitation depends on *R*). The solution (and the methods needed to obtain it) of the heterogeneous problem depends on the choice of the trait-dependent functions, which are detailed in below.

### Trait dependent trade-offs

We are interested in the cases where the proxy trait *x* provides the parasite with different levels of resource exploitation, and consequently to mortality due to possible life-history trade-offs. This trait will also provide a quantitative response to treatment, depending on the type of control strategy employed, so as to model different possible trait-specific treatments and their consequent heterogeneous efficacy. Therefore, in the following we will refer to exploitation, mortality and efficacy as functions of the proxy trait variable *x*. Our mathematical choices aim to capture the basic biological features of the trade-offs of interest, whilst maintaining mathematical tractability. Relaxation of such choices does not alter the qualitative features of our model, details are provided in the Supplemental material. References to the examples presented in the *Introduction* should be taken as qualitative connections underpinning our approach, rather than exact, detailed descriptions of the resistance, exploitation and mortality mechanisms at play.

### Exploitation

We assume exploitation rate *β* to be linearly affected by the trait variable *x*, that is:

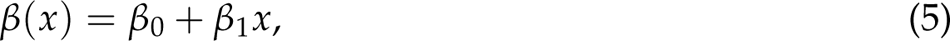

where *β*_0_ is the baseline rate, and *β*_1_ the trait-dependent contribution. We recall that in the micro-scenario, Eq. (5) corresponds to the transmission rate of the infection, and in the macro- scenario to the consumption rate.

### Mortality

In the microscenario, increased exploitation (*i*.*e*. transmission rate) is often associated with increased harm to the infected host (that is the Transmission-Virulence trade-off) (Nelson and May, 2020; Zhan et al., 2015; Laine and Barrès, 2013; Sacristán and Garćıa-Arenal, 2008; Montarry et al., 2006); in the macro- scenario, increased exploitation (*i*.*e*. consumption rate) is often associated with a reduced parasite lifespan due to, *e*.*g*., increased respiration or risk exposure (Strobbe et al., 2011; Stoks et al., 2005; Brodin and Johansson, 2004; Gotthard, 2000; Anholt et al., 2000; Werner and Anholt, 1993). In both scenarios, the respective trade-offs lead to an increase in parasite mortality. Accordingly, we assume that parasite mortality *δ^P^* can be linearly dependent on trait *x*:

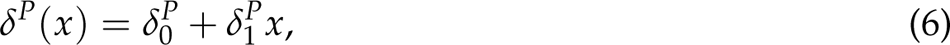

where the parameter *δ_0_^P^* is the baseline natural mortality, and *δ_1_^P^* is the trait-dependent contribution (Bolzoni and De Leo, 2013; Porco et al., 2005; Day and Proulx, 2004).

For values of both *δ_1_^P^* and *β*_1_ *>* 0, parasites with higher exploitation will also have higher mortality, consistent with the above trade-off hypothesis.

### Treatment efficacy

We assume that efficacy *ϕ*(*x*) is maximal for one of the extreme values of the trait (*x* = 0 and *x* = 1). Consistently with Porco et al. (2005), we choose a linear functional dependence:

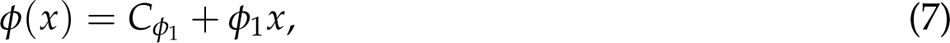

where the parameter *ϕ*_1_ represents the degree of correlation between the treatment and the trait (as explained in the next Subsection), and *C_ϕ_*_1_ is a normalisation factor. A graphical summary of the treatment spectrum is shown in Fig. 1 panel B (note that the bound *|ϕ*_1_*| <* 2 ensures a positive *ϕ*(*x*)). The linear assumption is made for the sake of mathematical tractability, although non-linear saturating choices are considered more realistic (Alizon, 2020). The effects of non-linearities can be explored semi-analytically, but they do not affect the quality of the results herein presented (details in the Supplemental Material).

**Figure 1:**
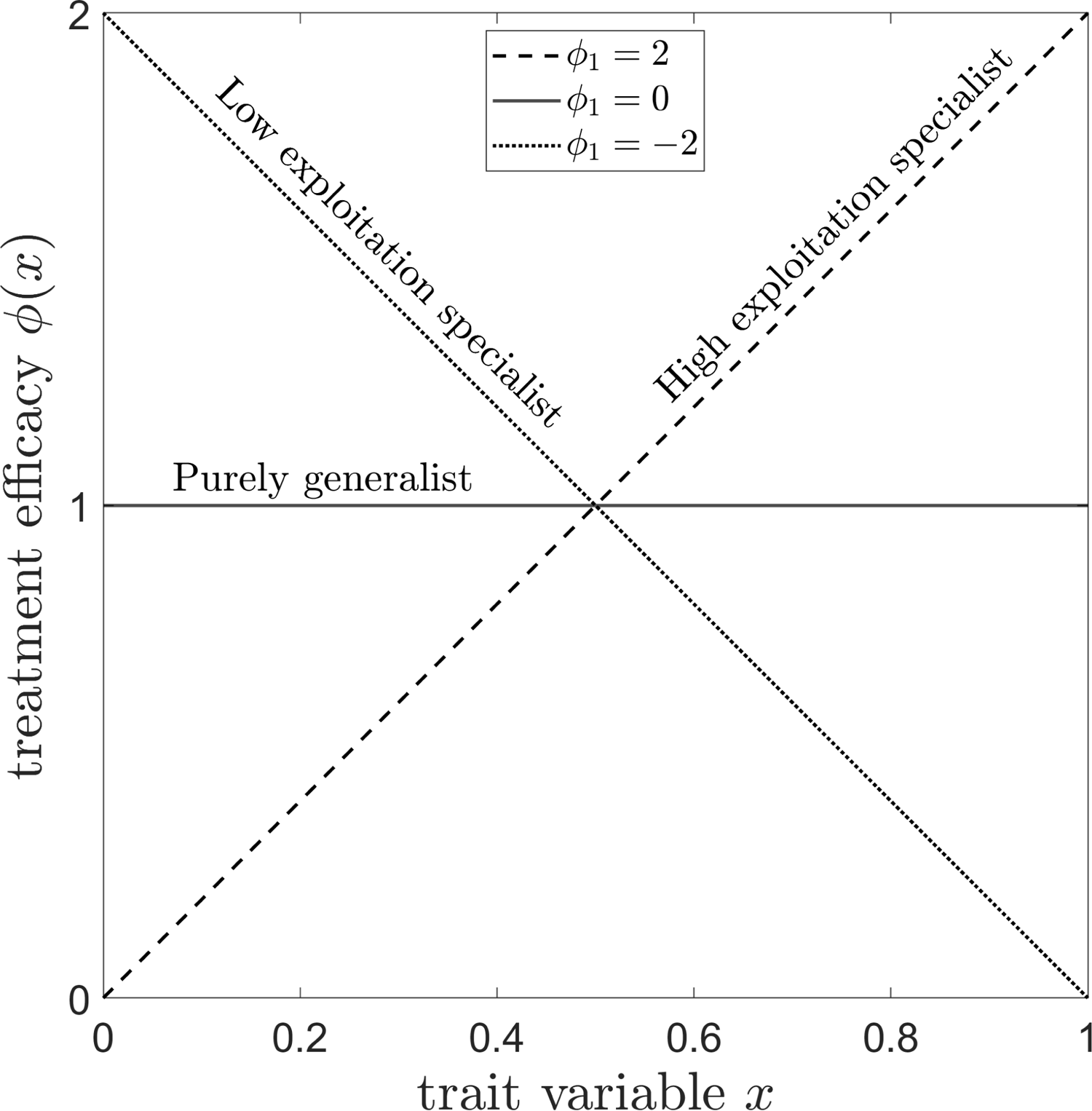
Spectrum of treatments with trait-dependent efficacy. The type of treatment is determined by the sign and the magnitude of the parameter *ϕ*_1_ *∈* [*−*2, 2]: positive *ϕ*_1_ models types with maximal effect on strains with higher exploitation levels (*x* = 1); negative *ϕ*_1_ models types with maximal effect on strains with lower exploitation levels (*x* = 0). Large *|ϕ*_1_*|* are specialised in targeting extreme values of the trait; small *|ϕ*_1_*|* are generalist types with more uniform action.

Combination of Eqns. (5), (6) and (7) captures the possible trade-offs occurring between trait-dependent treatment efficacy and the parasite’s exploitation levels. Ultimately, the ecoevolutionary dynamics of the system depends on the values of the parameters of the above functions, and thus on how a control strategy interacts with the proxy trait value *x*.

### Modelling treatment spectrum

We assume that the environmental and economic costs of the use of treatment are proportional to its application rate *γ*. We also assume that trait *x* determines the level of expression of a target trait, such as efflux pump expression, metabolic activity or proton motive force. The type of treatment then depends on how it correlates with the target trait, which is specified by the slope parameter *ϕ*_1_.

For instance, efflux pumps provide bacteria with resistance to chemical compounds, but make them vulnerable to attack by certain phages. Therefore, a standard antibiotic or pesticide treatment is more efficient on strains with lower levels of efflux pump expression (small *x*), and it is represented by a negative *ϕ*_1_; instead, a phage therapy benefits from higher levels of efflux pump expression (large *x*), and it is represented by a positive *ϕ*_1_.

Metabolic activity provides aphids an increased ability to consume and reproduce, but also makes them also more sensitive to pesticides, as they will tend to take up more toxic compounds. Therefore, the pesticide is more efficient on fast-exploiting strains (large *x*) and less efficient on slow-exploiting strains (small *x*); its slope *ϕ*_1_ is then positive; at the same time, reduced metabolic activity makes the aphids more vulnerable to natural enemies, so that a predator-based biocontrol will have a negative *ϕ*_1_.

Proton motive force reduces the import of aminoglycosides, but also the export of *β*-lactams. Therefore, aminoglycoside is more efficient against strains expressing less proton motive force (small *x*) and less efficient against strains with more proton motive force (large *x*); it is therefore represented by a negative *ϕ*_1_; however, *β*-lactam has the opposite effect, so it is represented by a positive *ϕ*_1_.

In any case, maximal efficacy is obtained at the extremes of the trait space (either *x* = 0 or *x* = 1), so it is assumed that the correlation of efficacy with traits is unimodal. In the absence of more precise data, we consider this assumption to be a reasonable starting point and leave other possible cases for discussion. The normalisation factor *C_ϕ_*_1_ = 1 *− ϕ*_1_/2 ensures that *ϕ*(*x*)d*x* is always normalised to 1, for any value of *ϕ*_1_. In addition to removing an arbitrary degree of freedom, this condition also imposes a plausible evolutionary constraint consistent with the notions of costs of resistance and of collateral sensitivity: resistance with respect to a particular treatment is paid for by high sensitivity with respect to others, as it has been observed in the examples illustrated above.

Recognising that a treatment may be either positively or negatively correlated with the proxy trait variable, we assume the existence of a continuous spectrum from which it is, in principle, possible to choose. This continuous spectrum mimics the possibility of synergistically combining different types of treatments, as it has been documented for phages-antibiotics (Kebriaei et al., 2022; Gu Liu et al., 2020), antibiotics-antivirulents (Rezzoagli et al., 2020), fungicide mixtures (van den Bosch et al., 2014), antibiotic mixtures (Nichol et al., 2019; Cokol et al., 2011), fungicide-biocontrol (Lima et al., 2006), pesticide-biocontrol (Foster et al., 2007). Crucially, we assume that the overall efficacy is the result of the sum of the single different treatment types. Therefore, any slope can be obtained by adjusting the relative proportions of antibiotic and phage therapy. Although we expect a full spectrum to be practically unavailable, it allows us to fully explore the eco-evolutionary behaviour of the system. Henceforth, with the term ‘specialist’ will refer to treatments with large degree of correlation (large *|ϕ*_1_*|*), and the term ‘generalist’ will refer to treatments with small degree of correlation (small *|ϕ*_1_*|*). A control strategy is thus characterised by one value of application rate *γ*, and one value of treatment type *ϕ*_1_.

### Agent-based numerical simulations

We compared the deterministic dynamics presented in our paper with numerical simulations of the equivalent agent-based dynamics. The Python code is available here. The agent-based dynamics simulates the stochastic events of *renewal*, *mortality*, *exploitation*, *treatment* and *mutation*, each occurring at a rate consistent with the deterministic Eqs. (2). The deterministic theory is expected to be consistent with our agent-based simulations as long as large populations are involved. Our aim is to provide a numerical validation of the existence and stability of the endemic equilibrium solution predicted by the theory. The analysis of finite size effects on the system is beyond the scope of this manuscript, although it is straightforward to perform once the agent-based codes are set up (Ardaševa et al., 2020). Details of the numerical simulations can be found in the Supplemental Material.

## Results

### Equilibrium trait distribution and evolutionary states

The state of the parasite population at equilibrium is described by the steady-state trait distribution 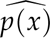, solution of:

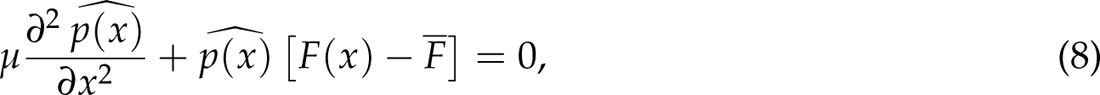

where the average quantities *β*, *δ^P^*, *ϕ*, appearing in *F*, are not known *a priori*, as they depend on the solution 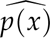 itself. Note that their time dependence has been dropped, since they reach a constant value at equilibrium. The above equation has a trivial solution 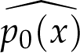 = 0, corresponding to the parasite-free equilibrium, and a non-trivial solution describing the endemic equilibrium, where resource and parasites coexist (mathematical details in the Supplemental Material).

The coexistence equilibrium exists (and is stable) as long as the following condition is satisfied:

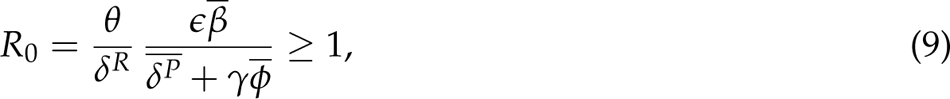

where *R*_0_ is the expected offspring produced by a parasite encountering an unexploited resource, per unit of biomass (also known as the basic reproduction number), for the heterogeneous system. Contrary to the classical formulation, the above condition cannot be calculated directly in terms of the ecological parameters, since it depends on average quantities that are not known *a priori*. Therefore, one must first solve Eq. (8) for a given set of parameters; then, calculate the average quantities and check with condition Eq. (9) the existence of the endemic equilibrium. In the Supplemental Material, we show that the behaviour of the solution to Eq. (8), with linear trait-dependent functions, is entirely determined by the following compound parameter Ω:

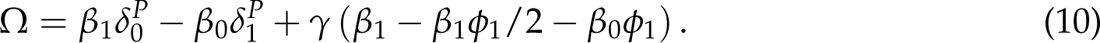

Particularly, if Ω < 0, the solution is monotonically decreasing and the trait distribution is mostly distributed close to the trait *x* = 0. We will refer to this as the *low-exploitation* state, because the corresponding trait has a minimum value for *x* = 0. On the other hand, if Ω *<ι><ιτ>></it>* 0, the solution is monotonically increasing and the trait distribution is mostly distributed close to the trait *x* = 1. Likewise, we will refer to this as the *high-exploitation* state.

Below, we characterise the phase diagram corresponding to a set of parameters, both in the presence and in absence of treatment. In this case, the proxy trait variable *x* simultaneously determines the levels of transmission and mortality of the parasite, as well as the efficacy of the treatment. In the absence of treatment (*i*.*e*. *γ* = 0), the parasite population can be found in either the *high* or *low exploitation* state, depending on the value of the ecological parameters. With reference to panel A of Fig. 2, *low* states will be favoured for large baseline exploitation *β*_0_ and trait-dependent mortality contribution *δ_1_^P^*; instead, *high* states will be favoured for large trait-dependent exploitation *β*_1_, and baseline mortality *δ_0_^P^*.

**Figure 2:**
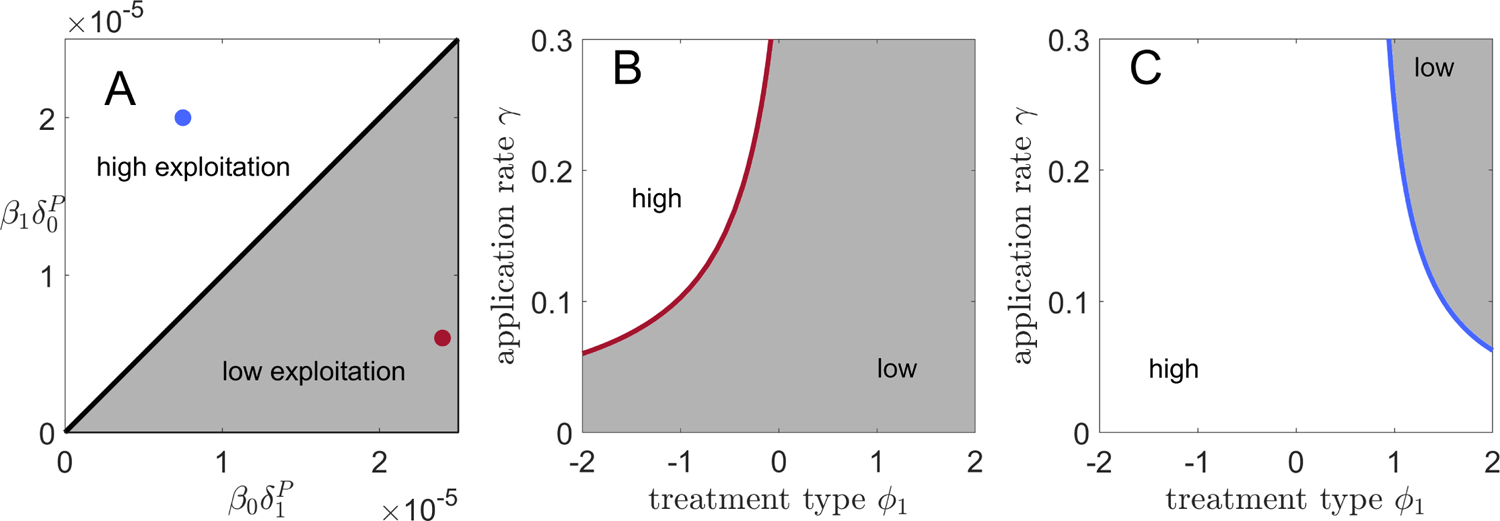
Trait distribution states. Panel A: state diagram of the trait distribution in absence of treatment. The parasite population can adapt towards either *low* (grey regions) or *high* (white regions) exploitation states. Panel B: a system initially in the *low* state (red point panel A) can switch towards the *high* state under a range of control strategies. The red curve separates the two regions of the control parameters. Panel C: likewise, a system initially in the *high* state (blue point panel A) can switch towards the *low* state under a range of control strategies. The blue curve separates the two regions of the control parameters. Parameters: *β*_0_ = 0.0001, *β*_1_ = *<u>^β^</u>*<u>^0^</u>, *δ^P^* = 0.12, *δ^P^* = 0.24 (for the red dot); *β*_0_ = *β*_1_ = 0.0001, *δ^P^* = 0.2, *δ^P^* = 0.075 (for the blue dot).

The introduction of treatment (*i*.*e*. *γ ̸*= 0) can lead to a change in state, depending on the control strategy (*ϕ*_1_, *γ*) employed. In panel B of Fig. 2, we show how a system initially in the *low* state (parameters corresponding to the red dot in panel A) adapts after treatment application, as a function of the control parameters *γ* and *ϕ*_1_. At low doses (*i*.*e*. low *γ*), the parasite population will remain in the *low* state, regardless of the type employed. However, increasing the application rate will eventually bring the system to the *high* state, if negative *ϕ*_1_ or generalist types are employed. If the system is initially in the *high* state in the absence of treatment (panel C of Fig. 2, blue dot of panel A), the complementary behaviour is observed.

In addition to affecting the evolutionary state of the system at endemic equilibrium, the control strategy also affects the amount of equilibrium resource. The adaptation towards the two possible states is shown in Fig. 3, where we plot the trajectories of the simulated agent-based dynamics (solid lines), differing for the treatment type employed. The other parameters (indicated in the figure caption) and initial conditions are identical. For *ϕ*_1_ = 2, the system adapts towards the *low* state, and towards the *high* state for *ϕ*_1_ = *−*2. Consequently, the resource reached at equilibrium is different in the two cases. The figure also shows the agreement between the analytical predictions for the equilibrium values (dashed lines), and the agent-based trajectories (solid lines), which holds for all sets of parameters considered in our analysis.

**Figure 3:**
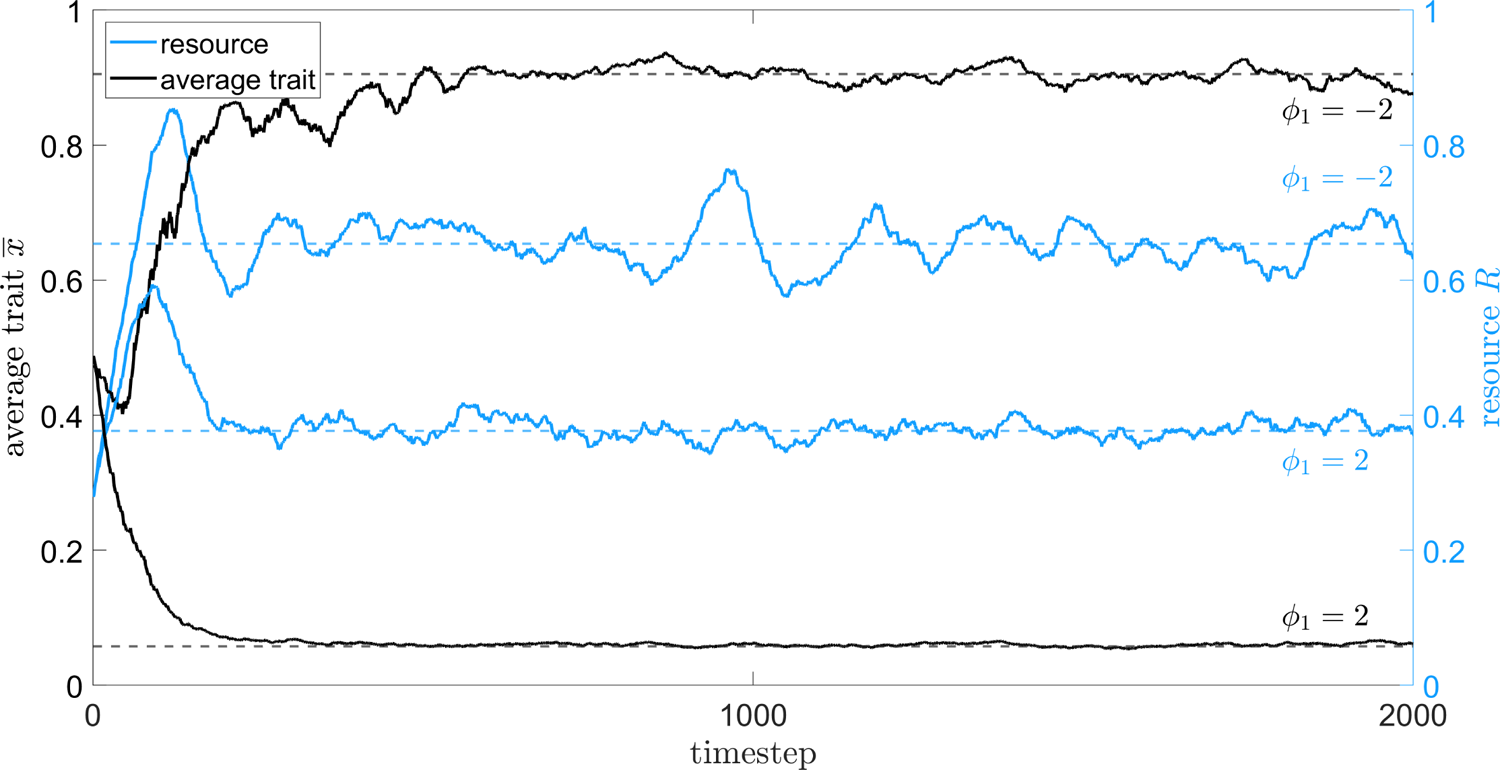
Simulated temporal trajectories. Solid lines: temporal trajectories of the resource *R* (rescaled with respect to *R*_0_) and of the average trait *x*, obtained from agent-based numerical simulations of the dynamics. Dashed lines: analytical equilibrium values predicted by the deterministic theory. The system is initialised with identical initial conditions and same application rate *γ* = 0.1, but different treatment type *ϕ*_1_. For *ϕ*_1_ = *±*2, resource and parasite average trait attain different equilibrium values. Other parameters: *θ* = 200, *δ_R_* = 0.04, *β*_0_ = 0.0001, *β*_1_ = *<u>^β^</u>*<u>^0^</u>, *δ^P^* = 0.12, *δ^P^* = 2*δ^P^*, *µ* = 7 *×* 10*^−^*^5^.

### Equilibrium resource and treatment effects

In the following, we will focus on the equilibrium resource 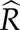, which can be considered as the amount of harvest with economic value (Vyska et al., 2016; Cunniffe et al., 2015). The population equilibria are obtained by setting Eqs. (2) to zero, and they are equivalent to the classical formulation. If *R*_0_ *<ι><ιτ><</it>* 1, we have a trivial parasite-free equilibrium:

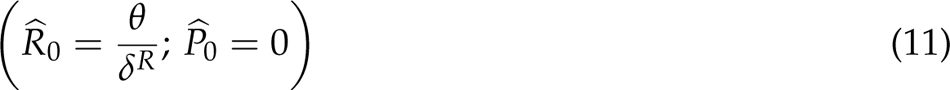

corresponding to the extinction of the parasite. If *R*_0_ *≥* 1, we have the following stable endemic equilibrium:

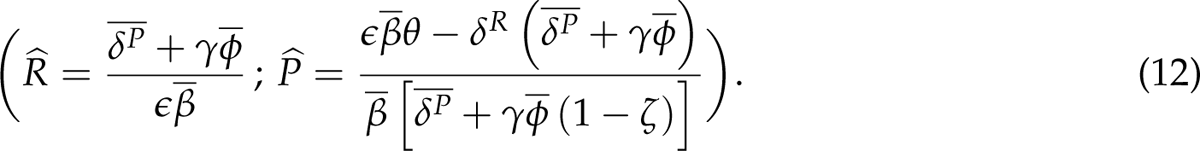

Note that the equilibrium averages are function of the control strategy (*ϕ*_1_, *γ*), since the equilibrium trait distribution 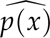 depends on such parameters. Therefore, Eq. (12) provides all the information about the complex relationship between resource production and control strategy, and it allows the systematic exploration of the whole parameter space. Note that the value of resource at the endemic equilibrium 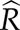 is independent of *ζ*. Therefore, a control strategy with such a quantity as objective will have the same outcome, regardless of the fate of the treated parasite. In the following, we will focus on a particular set of parameters as an example of the relevant behaviour of the system.

In Fig. 4 we plot the equilibrium resource 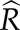 (rescaled with respect to the parasite-free resource 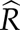_0_), as a function of application rate *γ*, and compare the effect of five different types of treatment: *ϕ*_1_ = *−*2, *−*1, 0, 1 and 2. We present two sets of parameters corresponding to the two opposite states of the parasite trait distribution, in the absence of treatment: the left panel corresponds to the red dot in panel A of Fig. 2, which is a low-exploitation state; the right panel corresponds to the blue dot in panel A of Fig. 2, which is a high-exploitation state. Note that employing a treatment type that is inconsistent with the state of the trait distribution in the absence of control (*e*.*g*. *ϕ*_1_ = 2 for the left panel, *ϕ*_1_ = *−*2 for the right panel), leads to a small increase in resource as the dose increases (green and blue curves, respectively). Instead, employing a type extremely specialised in the trait that dominates in the absence of control (*ϕ*_1_ = *−*2 for the left panel, *ϕ*_1_ = 2 for the right panel), leads to a more significant increase in resource, at least for low application rates (blue and green curves, respectively). However, as increasing the rate of application tends to push the system towards the opposite state, extremely specialised treatments can quickly become less effective, and the resource gain will eventually saturate. At this point, switching to a more moderate type (smaller *ϕ*_1_) rather than further increasing the application rate *γ*, will provide more resource gain. Depending on the value of the renewal rate *θ*, the system may eventually reach the parasite-free equilibrium (where 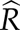 ≡ 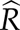_0_). Overall, when comparing the two panels, we find that the outcome of the control depends on the state of the parasite population in the absence of treatment, so that very efficient treatments in one case, may be very inefficient in the other. We also find that an increase in *γ*, regardless of the choice, always corresponds to an increase in the resource. Therefore, maximising the resource and minimising the treatment application are conflicting objectives. Nevertheless, it is possible to identify efficient strategies (*ϕ*_1_, *γ*), as explained below.

**Figure 4:**
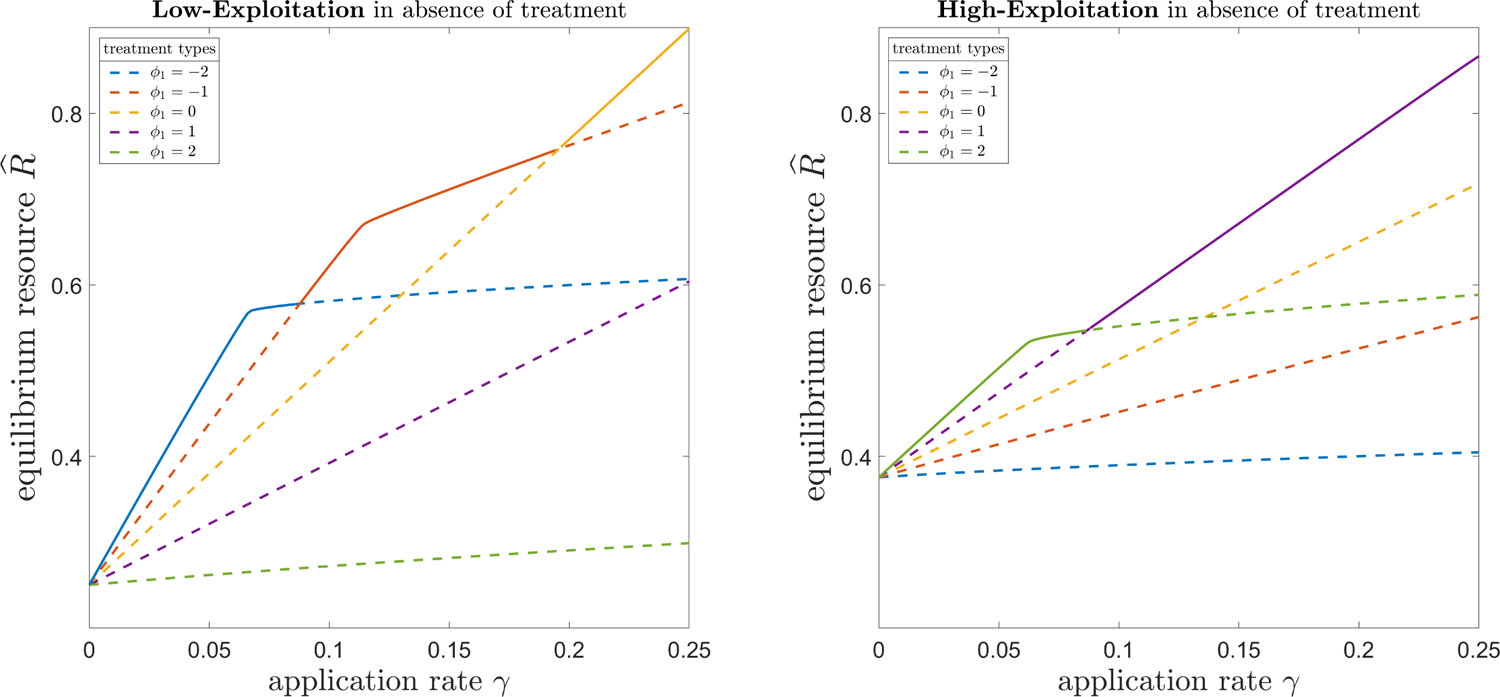
Equilibrium resource as function of control strategy. The resource at equilibrium *R* obtained using five different treatment types, as function of application rate *γ* (the y-axis is normalised with respect to the disease-free resource *R*_0_). Left panel: in absence of treatment, the system is in the low-exploitation state; parameters corresponding to red dot of Fig.2. Right panel: in absence of treatment, the system is in the high-exploitation state; parameters corresponding to blue dot of Fig. 2. Very specialised types (*ϕ*_1_ = *±*2) are efficient for low *γ*, but their correspondent gain in resource saturates as application rate is increased. Therefore, if application rate can be increased, more generalist treatment should be privileged. The Pareto-efficient strategies of the control strategy are highlighted with solid curves. Other parameters are: *µ* = 7 *×* 10*^−^*^5^, *θ* = 150, *δ_R_* = 0.04.

### Pareto-efficient strategies

Figure 4 shows that resource maximisation and treatment application minimisation are conflicting objectives. In the presence of conflicting objectives, multi-criteria analysis highlights the best compromises in the form of Pareto-efficient solutions (Kennedy et al., 2008). Among all the possible choices of our control strategy (*ϕ*_1_, *γ*), the Pareto-efficient solutions are those for which it is not possible to improve one objective without worsening the other. As such, they provide the decision maker with a smaller set of privileged alternatives to choose from, depending on the different management scenarios, and on the decision maker’s priorities.

The resulting Pareto-efficient solutions to our control strategy are identified by the solid curves in Fig. 4: choosing a control strategy (*ϕ*_1_, *γ*) different from the Pareto-efficient ones, will inevitably worsen the outcome (dashed curves), by either reducing the amount of resource or by increasing the costs associated with the treatment application.

We note that, when moving along the same type *ϕ*_1_, the resource shows a decrease in the incremental gain, for the following threshold application rate:

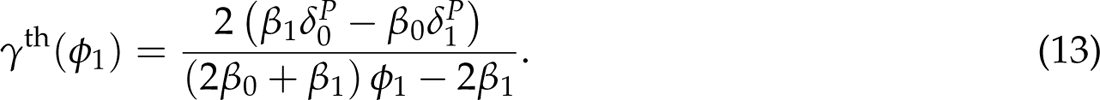

In the presence of a constraint on the application rate, the problem collapses to a unique objective function, which is optimised by (mathematical details in the Supplemental Material):

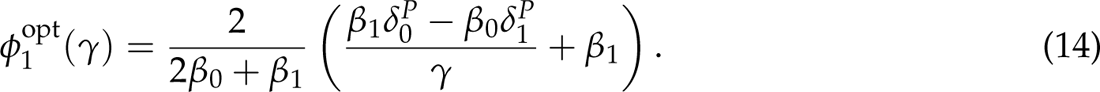

With respect to Fig. 2, the above equation corresponds to the curves separating the two states in the (*ϕ*_1_, *γ*) phase diagrams, for which Ω = 0. The effect of each parameter on *ϕ*^opt^(*γ*) is summarised in Table 2. Particularly, we note that: the optimal degree of specialisation is a decreasing function of application rate *γ*, so that extremely specialised treatments types will perform better at low *γ*; the optimal choice is independent of biomass conversion *ɛ*; exploitation and mortality rates (both baseline and trait-dependent contributions) play a non-trivial role in shaping the optimal choice. Overall, Eqs. (13) and (14) provides a qualitative insight into the role played of each ecological interaction in shaping the control strategy behaviour.

**Table 2:**
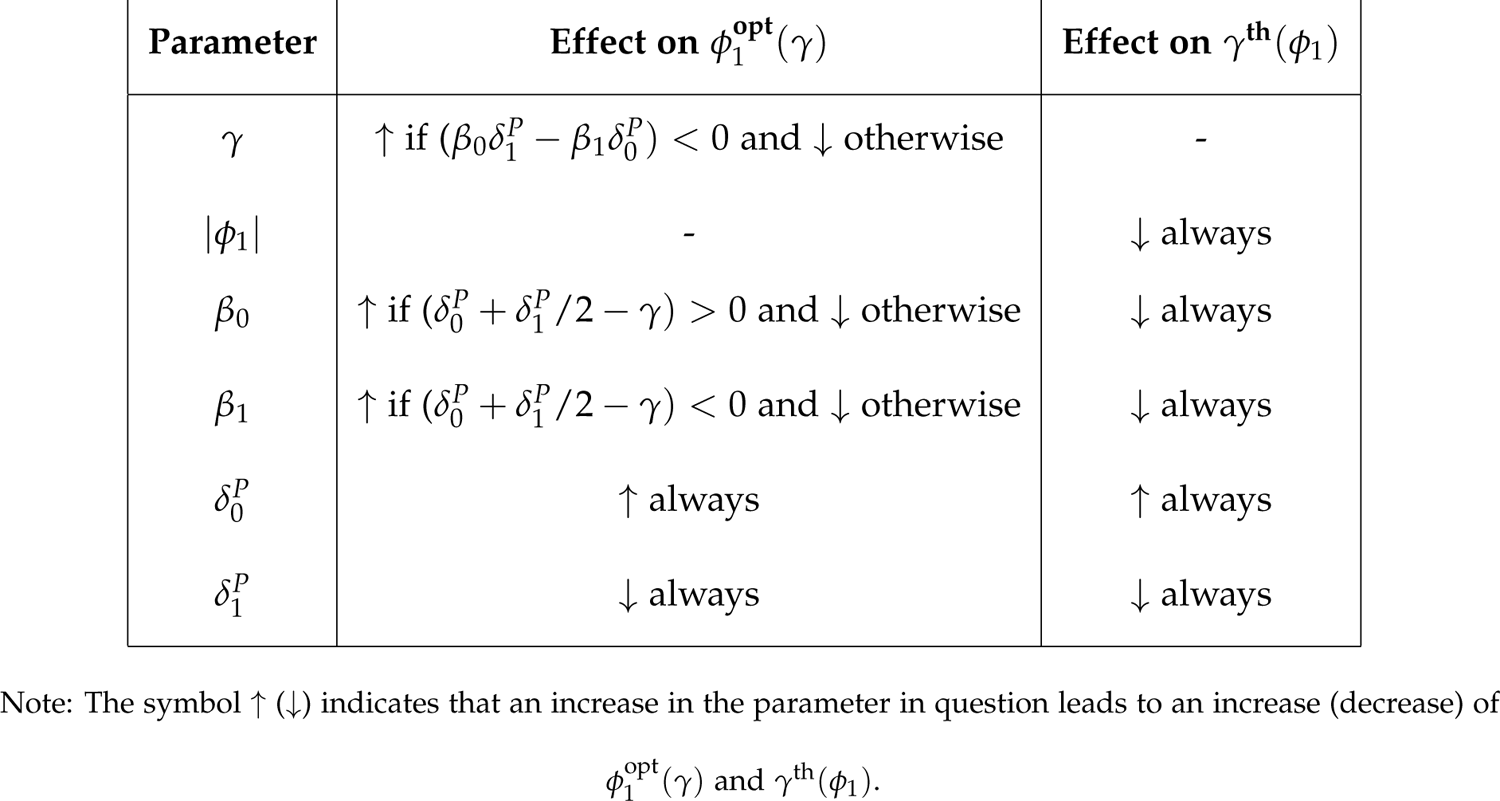
Effect of the parameters on the Pareto-efficient strategies.

## Discussion

We have developed a mathematical model to explore the implications of possible correlations between treatment efficacy and key traits of the parasite. We have considered a general parasite that may express continuous levels of exploitation and mortality (Eqs. 5 and 6), and a treatment that exerts an eradicant action, that may be either positively or negatively correlated with the above levels (Eq. 7 and Fig. 1), depending on the type of treatment. As a result of eco-evolutionary feedback, the parasite population can adapt towards evolutionary states dominated by either high or low exploitation levels (Fig. 2), and the final resource will depend on the control strategy employed (Fig. 4). The transition between these two possible states triggers several implications, depending on the management scenarios, which we discuss below.

Scenario 1): both application rate and treatment type are freely tunable; in this case, the efficient strategies are represented by a Pareto front (solid lines in Fig. 4). The Pareto front does not identify a single best strategy. Rather, it highlights a collection of best compromises between resource production and treatment use: whether to favour economic, environmental or ethical objectives will therefore depend on the priorities of the decision maker, as well as on how the resource and the application rate will map into real cost-benefit.

Scenario 2): the type of treatment may be constrained by the unavailability of alternatives or the inability to play with synergistic effects; in this case, there will be a threshold *γ*^th^, above which the decision maker should begin to question the benefit of further increasing the application rate. The economic impact of these saturation effects can be further assessed by including such information in economic evaluations of agricultural systems (Day et al., 2021; Ney et al., 2013; Paveley et al., 2001). Note that these saturation effects differ from those typically reported in the literature with treatment dose-response curves (Elderfield et al., 2018), which are accounted for by using non-linear, saturating functions. Here, saturation is due to a transition between opposite evolutionary states. This kind of saturation is then dynamic, and inherent to the eco-evolutionary nature of the system.

Scenario 3): the application rate may be constrained by limits on the use of antibiotics for safety reasons, or by limits on the spread of pesticides/copper/fungicides for legislative agricultural constraints. In this case, there will be a unique optimal type *ϕ*^opt^: as a general rule, very specialised treatment types should be employed at low application rates; instead, generalist types, with a more uniform action over the trait space, are likely to perform better at high application rates. This value represents the optimal choice from an ideal continuous spectrum of possibilities. We do not expect this spectrum to be fully available, or even possible to design in practice. Nvertheless, it should provide qualitative guidance to the decision maker when calibrating synergistic treatments.

The above scenarios assume that the control parameters *ϕ*_1_ and *γ* are independent, which may not always the case. However, a possible relation between application rate and treatment could be considered if the function *γ*(*ϕ*_1_) is known. Similarly, treatment application may be related to aspects of the host-parasite system, *e*.*g*. virulence or severity of the symptoms, (Porco et al., 2005). In such cases, it may be possible to derive optimal treatment conditions in terms of the epidemiological parameters of the parasite, provided that the corresponding functions are known.

Overall, our results can contribute to the ubiquitous call to reduce the use of chemicals in public health (Organization et al., 2014) and agriculture (Medina-Pastor and Triacchini, 2020). Specifically, they point towards many of the European Union’s principles (P) for sustainable farm management (Barzman et al., 2015): (P4) valuable synergistic use of alternative control methods, (P5) minimising environmental impact, (P6) reducing the use of chemicals, (P7) anti-resistance strategies. A concrete and urgent issue is the widespread use of copper in agriculture to combat plant diseases (Nunes et al., 2016). Due to its questionable efficacy and toxic side effects, there is an urgent call for its reduction (Tamm et al., 2018). In bacteria, resistance to heavy metals, such as copper, is mediated by efflux pumps, which are also involved in virulence to plants (Martinez et al., 2009; Ryan et al., 2007). Therefore, our theoretical framework could be used to support experimental studies of phage-copper synergy, which remain to be tested.

The assumption of a broad spectrum of treatment effects was motivated by the growing interest in developing therapies targeting specific traits, and the possibility of combining them synergistically with traditional synthetic treatments (Baym et al., 2016; Allen et al., 2014; Lima et al., 2006). So far, we have referred to collateral sensitivity, phage therapy and biocontrol, as examples of control strategies that interfere with heterogeneous traits involved in both resistance and resource exploitation. We argue that the phenomenon may be of interest to other treatments based on heterogeneity and evolutionary constraints, such as: anti-biofilm, photodynamic activation, and more on phage therapy.

Anti-biofilm and photodynamic activation rely on a planktonic-*vs*-sessile evolutionary trade-off (Feng et al., 2021; Tits et al., 2020; Almeida et al., 2014): some bacteria can either live and move individually (planktonic phase), or they can aggregate together into immobile structures, called biofilms (sessile phase). Targeting such structures is a desirable strategy, because they are involved in virulence and resistance to treatment. However, the efficacy of an anti-biofilm treatment would depend on the trait composition of the target bacterial population, which in turn depends on the trade-off between the two phases, similar to the phenomenon considered in this work.

A promising application of phage therapy is the use of phages that have bacterial capsules as receptors. Capsules are external polysaccharide layers that protect bacteria and facilitate attachment to host tissue. Capsules are therefore involved in host colonisation and in evasion of the immune response or treatment. Phage selection for reduced capsule production will impose trade-offs between virulence and antibiotic sensitivity (Song et al., 2021; Chiarelli et al., 2020).

A proper experimental investigation and measurement of specific trait-dependent effects would likely require two stages; first, a single-cell stage to detect heterogeneity in the level of expression of the trait of interest, and to measure the resulting trait-dependent interactions with the different treatments (Fernandes et al., 2011); this task could be performed using *in vitro* setups, such as microscopy, flow cytometry or of RNA-seq (Mohiuddin et al., 2020; Avraham et al., 2015). Second, a population stage where the overall effects on the demography and on the trait distributions can be monitored and measured; this task could be performed in controlled *in vivo* environments such as bioreactors (Levin and Udekwu, 2010), greenhouses, field plots and animal facilities (Band et al., 2016).

Our analysis has assumed a linear, monotonic and unimodal dose-response curve for treatment efficacy, so as to keep the model as simple as possible, and to favour mathematical tractability. In the Supplement Material, we show that the addition of non-linearities does not affect the qualitative behaviour of the system, as long as an evolutionary constraint is considered. Therefore, we conjecture that the emergence of optimal and threshold behaviours are inherent to the system, and that they are due to evolutionary constraints, rather than to its exact functional form. From a theoretical perspective, it would be interesting to prove this conjecture formally, for any kind of evolutionary constraint imposed on the treatment.

Instead, we expect that breaking the unimodal assumption will have non-trivial implications: on the one hand, a bimodal efficacy function would likely trigger multiple peaks in the trait distributions, therefore leading to possible branching phenomena where the parasite population splits into two separate subclasses of, *e*.*g*., mid-low and mid-high levels of exploitation; however, we are currently unaware of any evidence for bimodal treatment efficacy, and it would need to be motivated by (at least) phenomenological arguments. On the other hand, it is reasonable to imagine a treatment that would have maximal efficacy at an intermediate value of the proxy trait (rather than at the extremes), which would likely lead to non-monotonous trait distributions; accordingly, the variance of this putative bell-shaped efficacy could function as a tunable control parameter, governing possible intermediate scenarios between the homogeneous (rather flat, large variance) and the heterogeneous (rather peaked, small variance) extremes.

Although non-linear, concave-down functions are typically considered to relate exploitation and mortality, a simpler linear choice has allowed us to take full advantage of the mathematical analysis, whilst preserving the possibility to model exploitation-mortality trade-offs; it also provides the baseline results against which to compare non-linear functions, thus disentangling the role of non-linearities from the role of the trade-off alone. The relaxation to non-linear functions is discussed in the Supplement Material, where it is again shown that the qualitative behaviour of the linear case is preserved.

Although our analytical derivation of the optimal treatment type relies on many simplifying assumptions, our work highlights the qualitative role of the various epidemiological interactions, and it provides a starting point for introducing further elements of complexity. To conclude our discussion, we highlight some potentially interesting issues.

Although multidimensional trait spaces are rarely considered, the simultaneous presence of multiple traits encoding different features of the parasite would improve the realism of the model. Particularly, it would be interesting to explore the case where the treatment correlates with a subset of them, in order to mimic intervention policies with imperfect coverage (Walter and Lion, 2021). For instance, one could consider a pathogen endowed with a trait defining its transmission capacity, and a trait defining its disease-induced mortality; then, one could imagine the existence of an intervention affecting the transmission trait (*e*.*g*. protectant effects of a pesticide, quarantine policy, vaccination campaigns) and an intervention affecting the mortality trait. Performing a similar analysis, one might be able to compute the optimal combination of the two actions, and relate it to the geometry of the trait space (Miele et al., 2021), as well as to the possible evolutionary and economic constraints.

Our minimal model used simple demography for both resource and parasite dynamics. The introduction of more complex demographic functions (Cunniffe and Gilligan, 2010), could lead to to oscillating regimes around the endemic equilibrium. Such a maintained out-of-equilibrium demography results in a time-varying fitness landscape that could trigger out-of-equilibrium evolutionary responses, characterised by the alternation of low and high exploitation regimes of the trait distribution.

The expanding knowledge of the ecological, evolutionary and molecular interactions between parasites and treatments, coupled with the theoretical feedback should continue to provide opportunities to effectively address the challenge of disease management. Combining the practical development of trait-specific treatments with our theoretical methods of investigation (Saubin et al., 2021), may allow us to exploit heterogeneity of parasite populations - almost always seen as a key difficulty by allowing the evolution of resistance to human intervention - to our advantage. Ultimately, evolutionary epidemiology is an instantiation of a more general theory of evolutionary ecology (Lion, 2018). As such, the potential of the approach presented here can be exploited to investigate trait-dependent intervention in other domains, such as public health (Stearns, 2012) and cancer dynamics (Gatenby et al., 2009).

## Supporting information

Supplemental material with mathematical details and more analysis

## Literature Cited

Alcalde-Rico, M., Hernando-Amado, S., Blanco, P., and Martίnez, J. L. (2016). Multidrug efflux pumps at the crossroad between antibiotic resistance and bacterial virulence. Frontiers in microbiology, 7:1483.

Alekshun, M. N. and Levy, S. B. (2007). Molecular mechanisms of antibacterial multidrug resistance. Cell, 128(6):1037–1050.

Alizon, S. (2020). Treating symptomatic infections and the co-evolution of virulence and drug resistance. *bioRxiv*.

Alizon, S., Hurford, A., Mideo, N., and Van Baalen, M. (2009). Virulence evolution and the trade-off hypothesis: history, current state of affairs and the future. Journal of evolutionary biology, 22(2):245–259.

Alizon, S. and van Baalen, M. (2005). Emergence of a convex trade-off between transmission and virulence. The American Naturalist, 165(6):E155–E167.

Allen, R. C., Popat, R., Diggle, S. P., and Brown, S. P. (2014). Targeting virulence: can we make evolution-proof drugs? Nature Reviews Microbiology, 12(4):300–308.

Almeida, J., Tomé, J. P., Neves, M. G., Tomé, A. C., Cavaleiro, J. A., Cunha, Â., Costa, L., Faustino, M. A., and Almeida, A. (2014). Photodynamic inactivation of multidrug-resistant bacteria in hospital wastewaters: influence of residual antibiotics. Photochemical & Photobiological Sciences, 13(4):626–633.

Anderson, R. M. and May, R. M. (1979). Population biology of infectious diseases: Part i. Nature, 280(5721):361–367.

Andersson, D. I. and Hughes, D. (2010). Antibiotic resistance and its cost: is it possible to reverse resistance? Nature Reviews Microbiology, 8(4):260–271.

Anholt, B. R., Werner, E., and Skelly, D. K. (2000). Effect of food and predators on the activity of four larval ranid frogs. Ecology, 81(12):3509–3521.

Ardaševa, A., Anderson, A. R., Gatenby, R. A., Byrne, H. M., Maini, P. K., and Lorenzi, T. (2020). Comparative study between discrete and continuum models for the evolution of competing phenotype-structured cell populations in dynamical environments. Physical Review E, 102(4):042404.

Aulin, L., Liakopoulos, A., van der Graaf, P. H., Rozen, D. E., and van Hasselt, J. (2021). Design principles of collateral sensitivity-based dosing strategies. Nature communications, 12(1):1–14.

Avraham, R., Haseley, N., Brown, D., Penaranda, C., Jijon, H. B., Trombetta, J. J., Satija, R., Shalek, A. K., Xavier, R. J., Regev, A., et al. (2015). Pathogen cell-to-cell variability drives heterogeneity in host immune responses. Cell, 162(6):1309–1321.

Band, V. I., Crispell, E. K., Napier, B. A., Herrera, C. M., Tharp, G. K., Vavikolanu, K., Pohl, J., Read, T. D., Bosinger, S. E., Trent, M. S., et al. (2016). Antibiotic failure mediated by a resistant subpopulation in *Enterobacter cloacae*. Nature microbiology, 1(6):1–9.

Barbosa, C., Rö mhild, R., Rosenstiel, P., and Schulenburg, H. (2019). Evolutionary stability of collateral sensitivity to antibiotics in the model pathogen *Pseudomonas aeruginosa*. Elife, 8:e51481.

Bargués-Ribera, M. and Gokhale, C. S. (2020). Eco-evolutionary agriculture: Host-pathogen dynamics in crop rotations. PLoS computational biology, 16(1):e1007546.

Barzman, M., Bàrberi, P., Birch, A. N. E., Boonekamp, P., Dachbrodt-Saaydeh, S., Graf, B., Hom-mel, B., Jensen, J. E., Kiss, J., Kudsk, P., et al. (2015). Eight principles of integrated pest management. Agronomy for sustainable development, 35(4):1199–1215.

Bass, C., Puinean, A. M., Zimmer, C. T., Denholm, I., Field, L. M., Foster, S. P., Gutbrod, O., Nauen, R., Slater, R., and Williamson, M. S. (2014). The evolution of insecticide resistance in the peach potato aphid, *Myzus persicae*. Insect biochemistry and molecular biology, 51:41–51.

Baym, M., Stone, L. K., and Kishony, R. (2016). Multidrug evolutionary strategies to reverse antibiotic resistance. Science, 351(6268).

Beceiro, A., Tomás, M., and Bou, G. (2013). Antimicrobial resistance and virulence: a successful or deleterious association in the bacterial world? Clinical microbiology reviews, 26(2):185–230.

Blanquart, F., Lehtinen, S., Lipsitch, M., and Fraser, C. (2018). The evolution of antibiotic resistance in a structured host population. Journal of The Royal Society Interface, 15(143):20180040.

Bolzoni, L. and De Leo, G. A. (2013). Unexpected consequences of culling on the eradication of wildlife diseases: the role of virulence evolution. The American Naturalist, 181(3):301–313.

Boots, M. and Bowers, R. G. (2004). The evolution of resistance through costly acquired immunity. Proceedings of the Royal Society of London. Series B: Biological Sciences, 271(1540):715–723.

Boots, M. and Haraguchi, Y. (1999). The evolution of costly resistance in host-parasite systems. The american naturalist, 153(4):359–370.

Brodin, T. and Johansson, F. (2004). Conflicting selection pressures on the growth/predation-risk trade-off in a damselfly. Ecology, 85(11):2927–2932.

Bull, J. J. (1994). Virulence. Evolution, 48(5):1423–1437.

Bull, J. J., Vegge, C. S., Schmerer, M., Chaudhry, W. N., and Levin, B. R. (2014). Phenotypic resistance and the dynamics of bacterial escape from phage control. PloS one, 9(4):e94690.

Burmeister, A. R., Hansen, E., Cunningham, J. J., Rego, E. H., Turner, P. E., Weitz, J. S., and Hochberg, M. E. (2021). Fighting microbial pathogens by integrating host ecosystem interactions and evolution. Bioessays, 43(3):2000272.

Castle, M. D. and Gilligan, C. A. (2012). An epidemiological framework for modelling fungicide dynamics and control. PLoS One, 7(8):e40941.

Chan, B. K., Sistrom, M., Wertz, J. E., Kortright, K. E., Narayan, D., and Turner, P. E. (2016). Phage selection restores antibiotic sensitivity in mdr *Pseudomonas aeruginosa*. Scientific reports, 6(1):1–8.

Chan, B. K., Turner, P. E., Kim, S., Mojibian, H. R., Elefteriades, J. A., and Narayan, D. (2018). Phage treatment of an aortic graft infected with *Pseudomonas aeruginosa*. Evolution, medicine, and public health, 2018(1):60–66.

Chebotar, I. V., Emelyanova, M. A., Bocharova, J. A., Mayansky, N. A., Kopantseva, E. E., and Mikhailovich, V. M. (2021). The classification of bacterial survival strategies in the presence of antimicrobials. Microbial Pathogenesis, 155:104901.

Chiarelli, A., Cabanel, N., Rosinski-Chupin, I., Zongo, P. D., Naas, T., Bonnin, R. A., and Glaser, P. (2020). Diversity of mucoid to non-mucoid switch among carbapenemase-producing *Klebsiella pneumoniae*. BMC microbiology, 20(1):1–14.

Chisholm, R. H., Lorenzi, T., Desvillettes, L., and Hughes, B. D. (2016). Evolutionary dynamics of phenotype-structured populations: from individual-level mechanisms to population-level consequences. Zeitschrift fü r angewandte Mathematik und Physik, 67(4):100.

Cokol, M., Chua, H. N., Tasan, M., Mutlu, B., Weinstein, Z. B., Suzuki, Y., Nergiz, M. E., Costanzo, M., Baryshnikova, A., Giaever, G., et al. (2011). Systematic exploration of synergistic drug pairs. Molecular systems biology, 7(1):544.

consortium REX (2010). The skill and style to model the evolution of resistance to pesticides and drugs. Evolutionary Applications, 3(4):375–390.

Copin, R., Sause, W. E., Fulmer, Y., Balasubramanian, D., Dyzenhaus, S., Ahmed, J. M., Kumar, K., Lees, J., Stachel, A., Fisher, J. C., et al. (2019). Sequential evolution of virulence and resistance during clonal spread of community-acquired methicillin-resistant *Staphylococcus aureus*. Proceedings of the National Academy of Sciences, 116(5):1745–1754.

Corwin, J. A. and Kliebenstein, D. J. (2017). Quantitative resistance: more than just perception of a pathogen. The Plant Cell, 29(4):655–665.

Costerton, J. W., Stewart, P. S., and Greenberg, E. P. (1999). Bacterial biofilms: a common cause of persistent infections. Science, 284(5418):1318–1322.

Coyne, A. J. K., Stamper, K., Kebriaei, R., Holger, D. J., El Ghali, A., Morrisette, T., Biswas, B., Wilson, M., Deschenes, M. V., Canfield, G. S., et al. (2022). Phage cocktails with daptomycin and ampicillin eradicates biofilm-embedded multidrug-resistant *Enterococcus faecium* with preserved phage susceptibility. Antibiotics, 11(9):1175.

Cunniffe, N. J. and Gilligan, C. A. (2010). Invasion, persistence and control in epidemic models for plant pathogens: the effect of host demography. Journal of the Royal Society Interface, 7(44):439– 451.

Cunniffe, N. J., Koskella, B., Metcalf, C. J. E., Parnell, S., Gottwald, T. R., and Gilligan, C. A. (2015). Thirteen challenges in modelling plant diseases. Epidemics, 10:6–10.

Day, T. and Gandon, S. (2006). Insights from price’s equation into evolutionary. Disease evolution: models, concepts, and data analyses, 71:23.

Day, T., Kennedy, D. A., Read, A. F., and McAdams, D. (2021). The economics of managing evolution. PLoS biology, 19(11):e3001409.

Day, T., Parsons, T., Lambert, A., and Gandon, S. (2020). The price equation and evolutionary epidemiology. Philosophical Transactions of the Royal Society B, 375(1797):20190357.

Day, T. and Proulx, S. R. (2004). A general theory for the evolutionary dynamics of virulence. The American Naturalist, 163(4):E40–E63.

Dewachter, L., Fauvart, M., and Michiels, J. (2019). Bacterial heterogeneity and antibiotic survival: understanding and combatting persistence and heteroresistance. Molecular cell, 76(2):255–267.

Dutta, A., Croll, D., McDonald, B. A., and Barrett, L. G. (2020). Maintenance of variation in virulence and reproduction in populations of an agricultural plant pathogen. *bioRxiv*.

El Meouche, I. and Dunlop, M. J. (2018). Heterogeneity in efflux pump expression predisposes antibiotic-resistant cells to mutation. Science, 362(6415):686–690.

Elderfield, J. A., Lopez-Ruiz, F. J., van den Bosch, F., and Cunniffe, N. J. (2018). Using epidemiological principles to explain fungicide resistance management tactics: Why do mixtures outperform alternations? Phytopathology, 108(7):803–817.

Esteves, N. C. and Scharf, B. E. (2022). Flagellotropic bacteriophages: Opportunities and challenges for antimicrobial applications. International Journal of Molecular Sciences, 23(13):7084.

Fanning, S. and Mitchell, A. P. (2012). Fungal biofilms. PLoS pathogens, 8(4):e1002585.

Feng, Y., Tonon, C. C., Ashraf, S., and Hasan, T. (2021). Photodynamic and antibiotic therapy in combination against bacterial infections: efficacy, determinants, mechanisms, and future perspectives. Advanced Drug Delivery Reviews, 177:113941.

Fernandes, R. L., Nierychlo, M., Lundin, L., Pedersen, A. E., Tellez, P. P., Dutta, A., Carlquist, M., Bolic, A., Schäpper, D., Brunetti, A. C., et al. (2011). Experimental methods and modeling techniques for description of cell population heterogeneity. Biotechnology advances, 29(6):575– 599.

Forster, G. A. and Gilligan, C. A. (2007). Optimizing the control of disease infestations at the landscape scale. Proceedings of the National Academy of Sciences of the United States of America, 104(12):4984–4989.

Foster, S. P., Denholm, I., Poppy, G., Thompson, R., and Powell, W. (2011). Fitness trade-off in peach-potato aphids (*Myzus persicae*) between insecticide resistance and vulnerability to parasitoid attack at several spatial scales. Bulletin of Entomological Research, 101(6):659–666.

Foster, S. P., Tomiczek, M., Thompson, R., Denholm, I., Poppy, G., Kraaijeveld, A. R., and Powell, W. (2007). Behavioural side-effects of insecticide resistance in aphids increase their vulnerability to parasitoid attack. Animal Behaviour, 74(3):621–632.

Frank, S. A. (1996). Models of parasite virulence. The Quarterly review of biology, 71(1):37–78.

Furusawa, C., Horinouchi, T., and Maeda, T. (2018). Toward prediction and control of antibioticresistance evolution. Current opinion in biotechnology, 54:45–49.

Galvani, A. P. (2003). Epidemiology meets evolutionary ecology. Trends in Ecology & Evolution, 18(3):132–139.

Gandon, S., Mackinnon, M., Nee, S., and Read, A. (2003). Imperfect vaccination: some epidemiological and evolutionary consequences. Proceedings of the Royal Society of London. Series B: Biological Sciences, 270(1520):1129–1136.

Gandon, S., Mackinnon, M. J., Nee, S., and Read, A. F. (2001). Imperfect vaccines and the evolution of pathogen virulence. Nature, 414(6865):751–756.

Gatenby, R. A., Brown, J., and Vincent, T. (2009). Lessons from applied ecology: cancer control using an evolutionary double bind. Cancer research, 69(19):7499–7502.

Gefen, O. and Balaban, N. Q. (2009). The importance of being persistent: heterogeneity of bacterial populations under antibiotic stress. FEMS microbiology reviews, 33(4):704–717.

Gilligan, C. A. (2002). An epidemiological framework for disease management.

Gilligan, C. A. and van den Bosch, F. (2008). Epidemiological models for invasion and persistence of pathogens. Annu. Rev. Phytopathol., 46:385–418.

Giraud, E., Rychlik, I., and Cloeckaert, A. (2017). Antimicrobial resistance and virulence common mechanisms. Frontiers in microbiology, 8:310.

González, R., Butković, A., and Elena, S. F. (2019). Role of host genetic diversity for susceptibility-to-infection in the evolution of virulence of a plant virus. Virus evolution, 5(2):vez024.

Gotthard, K. (2000). Increased risk of predation as a cost of high growth rate: an experimental test in a butterfly: Predation as a cost of high growth rate. Journal of Animal Ecology, 69(5):896–902.

Gu Liu, C., Green, S. I., Min, L., Clark, J. R., Salazar, K. C., Terwilliger, A. L., Kaplan, H. B., Trautner, B. W., Ramig, R. F., and Maresso, A. W. (2020). Phage-antibiotic synergy is driven by a unique combination of antibacterial mechanism of action and stoichiometry. MBio, 11(4):e01462–20.

Gurney, J., Pradier, L., Griffin, J. S., Gougat-Barbera, C., Chan, B. K., Turner, P. E., Kaltz, O., and Hochberg, M. E. (2020). Phage steering of antibiotic-resistance evolution in the bacterial pathogen, *Pseudomonas aeruginosa*. Evolution, medicine, and public health, 2020(1):148–157.

Hall, R. J., Gubbins, S., and Gilligan, C. A. (2004). Invasion of drug and pesticide resistance is determined by a trade-off between treatment efficacy and relative fitness. Bulletin of Mathematical Biology, 66(4):825–840.

Hawkins, N. and Fraaije, B. (2018). Fitness penalties in the evolution of fungicide resistance. Annual review of phytopathology, 56:339–360.

Hethcote, H. W. (2000). The mathematics of infectious diseases. SIAM review, 42(4):599–653.

Hewitt, S. K., Foster, D. S., Dyer, P. S., and Avery, S. V. (2016). Phenotypic heterogeneity in fungi: importance and methodology. Fungal Biology Reviews, 30(4):176–184.

Kebriaei, R., Lev, K. L., Shah, R. M., Stamper, K. C., Holger, D. J., Morrisette, T., Kunz Coyne, A. J., Lehman, S. M., and Rybak, M. J. (2022). Eradication of biofilm-mediated methicillin-resistant *Staphylococcus aureus* infections in vitro: Bacteriophage-antibiotic combination. Microbiology Spectrum, 10(2):e00411–22.

Keeling, M. J. and Rohani, P. (2011). Modeling infectious diseases in humans and animals. Princeton university press.

Kennedy, M. C., Ford, E. D., Singleton, P., Finney, M., and Agee, J. K. (2008). Informed multiobjective decision-making in environmental management using pareto optimality. Journal of Applied Ecology, 45(1):181–192.

Kimura, M. (1965). A stochastic model concerning the maintenance of genetic variability in quantitative characters. Proceedings of the National Academy of Sciences of the United States of America, 54(3):731.

Korobeinikov, A. (2018). Immune response and within-host viral evolution: immune response can accelerate evolution. Journal of theoretical biology, 456:74–83.

Korobeinikov, A. and Wake, G. C. (2002). Lyapunov functions and global stability for sir, sirs, and sis epidemiological models. Applied Mathematics Letters, 15(8):955–960.

Lacey, L., Grzywacz, D., Shapiro-Ilan, D., Frutos, R., Brownbridge, M., and Goettel, M. (2015). Insect pathogens as biological control agents: back to the future. Journal of invertebrate pathology, 132:1–41.

Lafferty, K. D., DeLeo, G., Briggs, C. J., Dobson, A. P., Gross, T., and Kuris, A. M. (2015). A general consumer-resource population model. Science, 349(6250):854–857.

Laine, A.-L. and Barrès, B. (2013). Epidemiological and evolutionary consequences of life-history trade-offs in pathogens. Plant Pathology, 62:96–105.

Lässig, M., Mustonen, V., and Walczak, A. M. (2017). Predicting evolution. Nature ecology & evolution, 1(3):1–9.

Laure, N. N. and Ahn, J. (2022). Phage resistance-mediated trade-offs with antibiotic resistance in *Salmonella Typhimurium*. Microbial Pathogenesis, 171:105732.

Lázár, V., Martins, A., Spohn, R., Daruka, L., Grézal, G., Fekete, G., Számel, M., Jangir, P. K., Kintses, B., Csörgő, B., et al. (2018). Antibiotic-resistant bacteria show widespread collateral sensitivity to antimicrobial peptides. Nature microbiology, 3(6):718–731.

Lázár, V., Pal Singh, G., Spohn, R., Nagy, I., Horváth, B., Hrtyan, M., Busa-Fekete, R., Bogos, B., Méhi, O., Csörgő, B., et al. (2013). Bacterial evolution of antibiotic hypersensitivity. Molecular systems biology, 9(1):700.

Lehtinen, S., Blanquart, F., Lipsitch, M., Fraser, C., and with the Maela Pneumococcal Collaboration (2019). On the evolutionary ecology of multidrug resistance in bacteria. PLoS pathogens, 15(5):e1007763.

Lenski, R. E. (1998). Bacterial evolution and the cost of antibiotic resistance. Int Microbiol, 1(4):265– 270.

Levin, B. R. and Udekwu, K. I. (2010). Population dynamics of antibiotic treatment: a mathematical model and hypotheses for time-kill and continuous-culture experiments. Antimicrobial agents and chemotherapy, 54(8):3414–3426.

Lima, G., De Curtis, F., Piedimonte, D., Spina, A. M., and De Cicco, V. (2006). Integration of biocontrol yeast and thiabendazole protects stored apples from fungicide sensitive and resistant isolates of *Botrytis cinerea*. Postharvest biology and technology, 40(3):301–307.

Lion, S. (2018). Theoretical approaches in evolutionary ecology: environmental feedback as a unifying perspective. The American Naturalist, 191(1):21–44.

Lorenzi, T., Chisholm, R. H., and Clairambault, J. (2016). Tracking the evolution of cancer cell populations through the mathematical lens of phenotype-structured equations. Biology direct, 11(1):43.

Lyu, Z., Yang, A., Villanueva, P., Singh, A., and Ling, J. (2021). Heterogeneous flagellar expression in single salmonella cells promotes diversity in antibiotic tolerance. MBio, 12(5):e02374–21.

Maeda, T., Iwasawa, J., Kotani, H., Sakata, N., Kawada, M., Horinouchi, T., Sakai, A., Tanabe, K., and Furusawa, C. (2020). High-throughput laboratory evolution reveals evolutionary constraints in *Escherichia coli*. Nature communications, 11(1):1–13.

Maltas, J. and Wood, K. B. (2019). Pervasive and diverse collateral sensitivity profiles inform optimal strategies to limit antibiotic resistance. PLoS biology, 17(10):e3000515.

Mangalea, M. R. and Duerkop, B. A. (2020). Fitness trade-offs resulting from bacteriophage resistance potentiate synergistic antibacterial strategies. Infection and immunity, 88(7):e00926– 19.

Martinez, J. L., Sánchez, M. B., Martίnez-Solano, L., Hernandez, A., Garmendia, L., Fajardo, A., and Alvarez-Ortega, C. (2009). Functional role of bacterial multidrug efflux pumps in microbial natural ecosystems. FEMS microbiology reviews, 33(2):430–449.

Martίnez, S. R., Palacios, Y. B., Heredia, D. A., Agazzi, M. L., and Durantini, A. M. (2019). Phenotypic resistance in photodynamic inactivation unravelled at the single bacterium level. ACS Infectious Diseases, 5(9):1624–1633.

Mattei, M., Frunzo, L., D’acunto, B., Pechaud, Y., Pirozzi, F., and Esposito, G. (2018). Continuum and discrete approach in modeling biofilm development and structure: a review. Journal of mathematical biology, 76(4):945–1003.

McLeod, D. V. and Gandon, S. (2021). Understanding the evolution of multiple drug resistance in structured populations. Elife, 10:e65645.

Meaden, S., Paszkiewicz, K., and Koskella, B. (2015). The cost of phage resistance in a plant pathogenic bacterium is context-dependent. Evolution, 69(5):1321–1328.

Medina-Pastor, P. and Triacchini, G. (2020). The 2018 european union report on pesticide residues in food. EFSA Journal, 18(4):1–103.

Miele, L., Evans, R., and Azaele, S. (2021). Redundancy-selection trade-off in phenotype-structured populations. Journal of Theoretical Biology, 531:110884.

Mohiuddin, S. G., Kavousi, P., and Orman, M. A. (2020). Flow-cytometry analysis reveals persister resuscitation characteristics. BMC microbiology, 20(1):1–13.

Montarry, J., Corbiere, R., Lesueur, S., Glais, I., and Andrivon, D. (2006). Does selection by resistant hosts trigger local adaptation in plant–pathogen systems? Journal of Evolutionary Biology, 19(2):522–531.

Munita, J. M. and Arias, C. A. (2016). Mechanisms of antibiotic resistance. Microbiology spectrum, 4(2):4–2.

Nelson, P. and May, G. (2020). Defensive symbiosis and the evolution of virulence. The American Naturalist, 196(3):333–343.

Ney, B., Bancal, M.-O., Bancal, P., Bingham, I., Foulkes, J., Gouache, D., Paveley, N., and Smith, J. (2013). Crop architecture and crop tolerance to fungal diseases and insect herbivory. mechanisms to limit crop losses. European Journal of Plant Pathology, 135(3):561–580.

Nichol, D., Rutter, J., Bryant, C., Hujer, A. M., Lek, S., Adams, M. D., Jeavons, P., Anderson, A. R., Bonomo, R. A., and Scott, J. G. (2019). Antibiotic collateral sensitivity is contingent on the repeatability of evolution. Nature communications, 10(1):1–10.

Nunes, I., Jacquiod, S., Brejnrod, A., Holm, P. E., Johansen, A., Brandt, K. K., Priemé, A., and Sørensen, S. J. (2016). Coping with copper: legacy effect of copper on potential activity of soil bacteria following a century of exposure. FEMS microbiology ecology, 92(11).

Okusu, H., Ma, D., and Nikaido, H. (1996). Acrab efflux pump plays a major role in the antibiotic resistance phenotype of escherichia coli multiple-antibiotic-resistance (mar) mutants. Journal of bacteriology, 178(1):306–308.

Ons, L., Bylemans, D., Thevissen, K., and Cammue, B. P. (2020). Combining biocontrol agents with chemical fungicides for integrated plant fungal disease control. Microorganisms, 8(12):1930.

Organization, W. H. et al. (2014). Antimicrobial resistance global report on surveillance: 2014 summary. Technical report, World Health Organization.

Pál, C., Papp, B., and Lázár, V. (2015). Collateral sensitivity of antibiotic-resistant microbes. Trends in microbiology, 23(7):401–407.

Palmer, A. C. and Kishony, R. (2013). Understanding, predicting and manipulating the genotypic evolution of antibiotic resistance. Nature Reviews Genetics, 14(4):243–248.

Patyka, V., Buletsa, N., Pasichnyk, L., Zhitkevich, N., Kalinichenko, A., Gnatiuk, T., and Butsenko, L. (2016). Specifics of pesticides effects on the phytopathogenic bacteria. Ecological Chemistry and Engineering S, 23(2):311–331.

Paveley, N., Sylvester-Bradley, R., Scott, R., Craigon, J., and Day, W. (2001). Steps in predicting the relationship of yield on fungicide dose. Phytopathology, 91(7):708–716.

Perrier, A., Barlet, X., Rengel, D., Prior, P., Poussier, S., Genin, S., and Guidot, A. (2019). Spontaneous mutations in a regulatory gene induce phenotypic heterogeneity and adaptation of *Ralstonia solanacearum* to changing environments. Environmental microbiology, 21(8):3140–3152.

Perry, G. H. (2021). Evolutionary medicine. Elife, 10:e69398.

Pinheiro, F., Warsi, O., Andersson, D. I., and Lässig, M. (2021). Metabolic fitness landscapes predict the evolution of antibiotic resistance. Nature Ecology & Evolution, 5(5):677–687.

Porco, T. C., Lloyd-Smith, J. O., Gross, K. L., and Galvani, A. P. (2005). The effect of treatment on pathogen virulence. Journal of Theoretical Biology, 233(1):91–102.

Rezzoagli, C., Archetti, M., Mignot, I., Baumgartner, M., and Kü mmerli, R. (2020). Combining antibiotics with antivirulence compounds can have synergistic effects and reverse selection for antibiotic resistance in *Pseudomonas aeruginosa*. PLoS biology, 18(8):e3000805.

Rimbaud, L., Papäıx, J., Rey, J.-F., Barrett, L. G., and Thrall, P. H. (2018). Assessing the durability and efficiency of landscape-based strategies to deploy plant resistance to pathogens. PLoS computational biology, 14(4):e1006067.

Rodriguez-Gonzalez, R. A., Leung, C. Y., Chan, B. K., Turner, P. E., and Weitz, J. S. (2020). Quantitative models of phage-antibiotic combination therapy. MSystems, 5(1):e00756–19.

Roemhild, R. and Andersson, D. I. (2021). Mechanisms and therapeutic potential of collateral sensitivity to antibiotics. PLoS Pathogens, 17(1):e1009172.

Ryan, R. P., Ryan, D. J., Sun, Y.-C., Li, F.-M., Wang, Y., and Dowling, D. N. (2007). An acquired efflux system is responsible for copper resistance in *Xanthomonas* strain ig-8 isolated from china. FEMS microbiology letters, 268(1):40–46.

Sacristán, S. and Garćıa-Arenal, F. (2008). The evolution of virulence and pathogenicity in plant pathogen populations. Molecular plant pathology, 9(3):369–384.

Sasaki, A., Lion, S., and Boots, M. (2021). Antigenic escape selects for the evolution of higher pathogen transmission and virulence. Nature ecology & evolution, pages 1–12.

Saubin, M., Louet, C., Bousset, L., Fabre, F., Fudal, I., Grognard, F., Mailleret, L., Stoeckel, S., Touzeau, S., Petre, B., et al. (2021). Improving the design of sustainable crop protection strategies thanks to population genetics concepts.

Schrö ter, L. and Dersch, P. (2019). Phenotypic diversification of microbial pathogens—cooperating and preparing for the future. Journal of molecular biology, 431(23):4645– 4655.

Schuster, P. and Sigmund, K. (1983). Replicator dynamics. Journal of theoretical biology, 100(3):533– 538.

Shoval, O., Sheftel, H., Shinar, G., Hart, Y., Ramote, O., Mayo, A., Dekel, E., Kavanagh, K., and Alon, U. (2012). Evolutionary trade-offs, pareto optimality, and the geometry of phenotype space. Science, 336(6085):1157–1160.

Song, L., Yang, X., Huang, J., Zhu, X., Han, G., Wan, Y., Xu, Y., Luan, G., and Jia, X. (2021). Phage selective pressure reduces virulence of hypervirulent Klebsiella pneumoniae through mutation of the wzc gene. Frontiers in microbiology, 12.

Stearns, S. C. (2012). Evolutionary medicine: its scope, interest and potential. Proceedings of the Royal Society B: Biological Sciences, 279(1746):4305–4321.

Stoks, R., Block, M. D., Van De Meutter, F., and Johansson, F. (2005). Predation cost of rapid growth: behavioural coupling and physiological decoupling. Journal of Animal Ecology, 74(4):708–715.

Strobbe, F., McPeek, M. A., De Block, M., and Stoks, R. (2011). Fish predation selects for reduced foraging activity. Behavioral Ecology and Sociobiology, 65(2):241–247.

Suzuki, S., Horinouchi, T., and Furusawa, C. (2014). Prediction of antibiotic resistance by gene expression profiles. Nature communications, 5(1):1–12.

Taber, H. W., Mueller, J. P., Miller, P. F., and Arrow, A. (1987). Bacterial uptake of aminoglycoside antibiotics. Microbiological reviews, 51(4):439–457.

Tamm, L., Pertot, I., Schmitt, A., Verrastro, V., Magid, J., Bü nemann, E., Mö ller, K., Athanasi-adou, S., Experton, C., Leiber, F., et al. (2018). Replacement of contentious inputs in organic farming systems (relacs)–a comprehensive horizon 2020 project. In Book of Abstracts. 6th International Conference on Organic Agriculture Sciences (ICOAS), 7-9 November 2018, Eisenstadt, Austria, page 47.

Taylor, N. and Cunniffe, N. (2022). Modelling quantitative fungicide resistance and breakdown of resistant cultivars: designing integrated disease management strategies for Septoria of winter wheat. *bioRxiv*.

Tits, J., Cammue, B. P., and Thevissen, K. (2020). Combination therapy to treat fungal biofilm-based infections. International Journal of Molecular Sciences, 21(22):8873.

Torres-Barceló, C. and Hochberg, M. E. (2016). Evolutionary rationale for phages as complements of antibiotics. Trends in microbiology, 24(4):249–256.

van den Bosch, F. and Gilligan, C. A. (2008). Models of fungicide resistance dynamics. Annu. Rev. Phytopathol., 46:123–147.

Van Den Bosch, F. and Gilligan, C. A. (2008). Models of fungicide resistance dynamics. Annual Review of Phytopathology, 46:123–147.

van den Bosch, F., Paveley, N., van den Berg, F., Hobbelen, P., and Oliver, R. (2014). Mixtures as a fungicide resistance management tactic. Phytopathology, 104(12):1264–1273.

Van Emden, H. F. and Harrington, R. (2017). Aphids as crop pests. Cabi.

Vincent, B. M., Lancaster, A. K., Scherz-Shouval, R., Whitesell, L., and Lindquist, S. (2013). Fitness trade-offs restrict the evolution of resistance to amphotericin b. PLoS biology, 11(10):e1001692.

Vyska, M., Cunniffe, N., and Gilligan, C. (2016). Trade-off between disease resistance and crop yield: a landscape-scale mathematical modelling perspective. Journal of the Royal Society Interface, 13(123):20160451.

Walter, A. and Lion, S. (2021). Epidemiological and evolutionary consequences of periodicity in treatment coverage. Proceedings of the Royal Society B, 288(1946):20203007.

Weigel, W. and Dersch, P. (2018). Phenotypic heterogeneity: a bacterial virulence strategy. Microbes and infection, 20(9-10):570–577.

Weiße, A. Y., Oyarzú n, D. A., Danos, V., and Swain, P. S. (2015). Mechanistic links between cellular trade-offs, gene expression, and growth. Proceedings of the National Academy of Sciences, 112(9):E1038–E1047.

Werner, E. and Anholt, B. (1993). Ecological consequences of the trade-off between growth and mortality rates mediated by foraging activity. The American naturalist, 142(2):242–272.

Zhan, J., Thrall, P. H., Papäıx, J., Xie, L., and Burdon, J. J. (2015). Playing on a pathogen’s weakness: using evolution to guide sustainable plant disease control strategies. Annual review of phytopathology, 53:19–43.

Zurita-Gutiérrez, Y. H. and Lion, S. (2015). Spatial structure, host heterogeneity and parasite virulence: implications for vaccine-driven evolution. Ecology letters, 18(8):779–789.

