## Supplemental material with mathematical details and more analysis for "Evolutionary epidemiology consequences of trait-dependent control of heterogeneous parasites"

### Supplemental Material for "**Evolutionary epidemiology consequences of trait-dependent control of heterogeneous parasites**"

#### 1 From the $RP$ to the $RPx$ formulation

In this section, we shall derive the heterogeneous formulation of the $RPx$ model presented in the main text, from the classical $RP$ formulation. We recall that the starting single-strain model is:

$$\begin{aligned} \frac{dR}{dt} &= \theta - \delta_R - \beta RP + \zeta\gamma\phi P \\ \frac{dP}{dt} &= \epsilon\beta RP - \big(\delta_P + \nu + \gamma\phi\big)P, \end{aligned} \tag{2}$$

where $R$ and $P$ are, respectively, the resource and parasaite biomass.

Trait heterogeneity in the parasite is accounted for by introducing the trait variable $x \in \mathcal{T} = [0;1]$, determining the levels of exploitation $\beta(x)$, additional mortality $\delta^P(x)$ and treatment efficacy $\phi(x)$. Denoting with $P(x)$ the amount of parasite biomass with trait $x$, the dynamics of the heterogeneous system is given by:

$$\frac{dR}{dt} = \theta - \delta_R R - \overline{\beta(t)} RP + \zeta\gamma\phi P \tag{3}$$

$$\frac{dP}{dt} = \epsilon\overline{\beta(t)} RP - \left[\overline{\delta_P(t)} + \gamma\overline{\phi(t)}\right] P \tag{4}$$

$$\frac{dP(x)}{dt} = \mu\frac{\partial^2 P(x)}{\partial x^2} + \epsilon\beta(x)RP(x) - \left[\delta^P(x) + \gamma\phi(x)\right] P(x), \tag{5}$$

where the bar notation indicates the average over the infected population, *e.g.*:

$$\overline{\beta(t)} = \int_{\mathcal{T}} \beta(x)\frac{P(x)}{P}\, dx. \tag{6}$$

In order to highlight the evolutionary nature of the dynamics, we derive an equivalent formulation that makes use of the trait distribution $p(x)$ defined as:

$$p(x) = \frac{P(x)}{P}. \tag{7}$$

The trait distribution determines the density of parasites with trait $x$. As such, this quantity is normalised:

$$\int_{\mathcal{T}} p(x)\, dx = \int_{\mathcal{T}} \frac{P(x)}{P}\, dx = \frac{P}{P} = 1. \tag{8}$$

The equation governing the temporal evolution of the trait distribution is obtained by taking the time derivative of its definition Eq. 7, and applying the chain rule:

$$\frac{d\,p(x)}{dt} = \frac{d}{dt}\frac{P(x)}{P} = \frac{1}{P}\frac{dP(x)}{dt} - \frac{P(x)}{P^2}\frac{dP}{dt} \tag{9}$$

$$= \mu\frac{\partial^2 \frac{P(x)}{P}}{\partial x^2} + \epsilon\beta(x)R\frac{P(x)}{P} - \left[\delta + \nu(x) + \gamma\phi(x)\right]\frac{P(x)}{P} - \frac{P(x)}{P}\left\{\epsilon\overline{\beta(t)}R - \left[\overline{\delta^P(t)} + \gamma\overline{\phi(t)}\right]\right\}, \tag{10}$$

where in the second line we have used Eqs. 4-5. Using Eq. 7 and performing simple algebraic steps, we finally obtain:

$$\frac{d\,p(x)}{dt} = \mu\frac{\partial^2\,p(x)}{\partial x^2} + p(x)\left[F(x) - \overline{F(t)}\right], \tag{11}$$

with:

$$F(x) = \epsilon\beta(x)R - \delta^P(x) - \gamma\phi(x), \tag{12}$$

as presented in the main text. Equation 11 is a non-linear integro-differential equation known in the theoretical literature as the Replicator-Mutator Equation (RME). The RME is closed with the following conditions:

$$\left.\begin{aligned} \int_{\mathcal{T}} p(x)\,dx &= 1 \\ \frac{\partial\,p(x)}{\partial\,x}\Big|_{x=0} = \frac{\partial\,p(x)}{\partial\,x}\Big|_{x=1} &= 0 \end{aligned}\right\} \quad \forall\, t, \tag{13}$$

corresponding to the two physical constraints satisfied by the trait distribution: normalisation at every time; and zero flux across the boundary of the strain domain. The former condition holds by definition (Eq. 8), whereas the latter derives from the reflecting nature of mutations close to the boundary (mutations leading to values of $x$ outside $\mathcal{T}$ are rejected).

### 2 Endemic equilibrium

#### Condition of existence

The resource and parasite biomass at endemic equilibrium are given by:

$$\left(\widehat{R} = \frac{\overline{\delta^P} + \gamma\overline{\phi}}{\epsilon\overline{\beta}}\ ;\ \widehat{P} = \frac{\epsilon\overline{\beta}\theta - \delta^R\left(\overline{\delta^P} + \gamma\overline{\phi}\right)}{\overline{\beta}\left[\overline{\delta^P} + \gamma\overline{\phi}\left(1 - \zeta\right)\right]}\right). \tag{14}$$

The endemic equilibrium has biological meaning if $\widehat{P} > 0$, that is if:

$$\epsilon\theta\overline{\beta} - \delta^R\left(\overline{\delta^P} + \overline{\phi}\right) > 0. \tag{15}$$

Dividing the above condition by the quantity $\epsilon\theta\overline{\beta} > 0$ and rearranging terms, we obtain:

$$R_0 = \frac{\theta}{\delta_R}\frac{\epsilon\overline{\beta}}{\overline{\delta^P} + \gamma\overline{\phi}} > 1, \tag{16}$$

consistently with the main text. The equilibrium average quantities $\overline{\beta}, \overline{\delta^P}, \overline{\phi}$ are determined by the equilibrium trait distribution $\widehat{p(x)}$, solution of:

$$\mu\frac{\partial^2\widehat{p(x)}}{\partial x^2} + \widehat{p(x)}\left\{\epsilon\beta(x)\widehat{R} - \delta^P(x) - \gamma\phi(x) - \left[\epsilon\overline{\beta}\widehat{R} - \overline{\delta^P} - \gamma\overline{\phi}\right]\right\} = 0. \tag{17}$$

The solution depending on its average quantities, Eq. 17 can be solved employing self-consistent methods, as explained in the following.

#### Analytical solution of the trait distribution

Under the choice of linear trait-dependant epidemiological trait functions (Eqs. 5 – 7 of the main text), Eq. 17 is:

$$\mu \frac{\partial^2 \widehat{p(x)}}{\partial x^2} + \frac{\Omega}{\beta_0 + \beta_1 \overline{x}} \widehat{p(x)} \big( x - \overline{x} \big) = 0, \tag{18}$$

with:

$$\Omega = \beta_1 \delta_0^P - \beta_0 \delta_1^P + \gamma \big( \beta_1 - \beta_1 \phi_1 / 2 - \beta_0 \phi_1 \big). \tag{19}$$

If $\Omega > 0$, performing the transformation of variable $z = \sqrt[3]{\frac{\Omega}{\mu[\beta_0 + \beta_1 x]}} \, (\overline{x} - x)$, Eq. 18 becomes:

$$\frac{d^2 p(z)}{d z^2} - z \, p(z) = 0, \tag{20}$$

which is the well known Airy differential equation (1). The solution is then a linear combination of the Airy functions of first and second kind which, back in the original variable, reads:

$$\widehat{p_{\overline{x}}(x)} = \mathcal{Z} \Big\{ \mathrm{Ai} \left[ \gamma \left( \overline{x} - x \right) \right] + \mathcal{C} \, \mathrm{Bi} \left[ \gamma \left( \overline{x} - x \right) \right] \Big\}, \quad \gamma = \sqrt[3]{\frac{|\Omega|}{\mu \left( \beta_0 + \beta_1 \overline{x} \right)}}, \tag{21}$$

with $\mathcal{Z}$ and $\mathcal{C}$ constants of integration given by (prime notation indicates the $x$ derivative):

$$\begin{cases} \mathcal{C} = -\frac{\mathrm{Ai}'(\gamma\overline{x})}{\mathrm{Bi}'(\gamma\overline{x})} \\ \mathcal{Z}^{-1} = \int_{\mathcal{T}} \mathrm{Ai}\left[\gamma\left(\overline{x} - x\right)\right] + \mathcal{C}\,\mathrm{Bi}\left[\gamma\left(\overline{x} - x\right)\right]\, dx \end{cases} \tag{22}$$

Equation 22 represents the $\overline{x}$-family of solutions to Eq. 18. The correct solution can be identified self-consistently by solving the constraint related to the average quantity:

$$\overline{x} = \int_{\mathcal{T}} x\, \widehat{p_{\overline{x}}(x)}\, dx. \tag{23}$$

This last step can be performed numerically by means of standard routines such as MATLAB or Python's *fsolve*() function. If $\Omega < 0$, equivalent solutions to Eq. 22 can be found upon replacing $x$ with $(1 - x)$.

### Optimal treatment type

At endemic equilibrium, the average mortality, exploitation and treatment efficacy, are respectively:

$$\begin{cases} \overline{\delta^P} = \delta_0^P + \delta_1^P \overline{x} \\ \overline{\beta} = \beta_0 + \beta_1 \overline{x} \\ \overline{\phi} = \phi_1 \overline{x} + 1 - \phi_1/2. \end{cases} \tag{24}$$

Inserting the above equations into the endemic equilibrium resource $\widehat{R}$ Eq. 14, we get:

$$\widehat{R} = \frac{\delta_0^P + \delta_1^P \overline{x} + \gamma \phi_1 \overline{x} + \gamma \left(1 - \phi_1/2\right)}{\beta_0 + \beta_1 \overline{x}}. \tag{25}$$

For a given set of the parameters $\delta_0^P, \delta_1^P, \beta_0, \beta_1$ and $\gamma$, the treatment type $\phi_1$ that maximises the equilibrium resource satisfies:

$$\frac{\partial \widehat{R}}{\partial \phi_1} = 0 \quad , \quad \frac{\partial^2 \widehat{R}}{\partial \phi_1^2} < 0. \tag{26}$$

Performing the first derivative with respect to $\phi_1$ of $\widehat{R}$, we get:

$$\frac{\partial \widehat{R}}{\partial \phi_1} = \frac{\left(\delta_1^P \overline{x}_{\phi_1} + \gamma \overline{x} + \gamma \phi_1 \overline{x}_{\phi_1} - \gamma/2\right)\left(\beta_0 + \beta_1 \overline{x}\right) - \left(\delta_0^P + \delta_1^P \overline{x} + \gamma \phi_1 \overline{x} + \gamma - \gamma \phi_1/2\right)\beta_1 \overline{x}_{\phi_1}}{\epsilon\left(\beta_0^2 + \beta_1 \overline{x}\right)^2}, \tag{27}$$

where the subscript notation indicates the derivative with respect to $\phi_1$. Arranging terms (and neglecting the denominator which is always positive), we get:

$$\frac{\partial \widehat{R}}{\partial \phi_1} = 0 \Longleftrightarrow \gamma\left(\beta_0 + \beta_1 \overline{x}\right)\left(\overline{x} - 1/2\right) - \Omega\, \overline{x}_{\phi_1} = 0. \tag{28}$$

The above condition is readily solved by observing that $\Omega = 0$ for $\overline{x} = 1/2$. Hence, the candidate for the optimal type is:

$$\phi_1^{\text{opt}} \quad \text{such that} \quad \overline{x} = \frac{1}{2}. \tag{29}$$

Checking the condition on the second derivative for the above candidate is straightforward. By setting the definition of $\Omega$ Eq. 19 to zero, we finally obtain the value of the optimal type $\phi_1^{\rm opt}(\gamma)$ as function of treatment application rate (as well as of the other parameters):

$$\phi_1^{\rm opt}(\gamma) = \frac{2}{2\beta_0 + \beta_1}\left[\frac{\beta_1\delta_0^P - \beta_0\delta_1^P}{\gamma} + \beta_1\right], \tag{30}$$

as reported in the main text.

### 3 Agent-based numerical simulations

The results predicted by our deterministic theory are checked with numerical simulation of the agent-based dynamics. The python codes to reproduce the agent-based simulations are available here.

The system is composed of discrete units of resource and parasites. The former is represented by the scalar number $R$ of total resource units in the population. The latter is represented by an array of dimension $P$, where $P$ is the total number parasite units. Each element $P$ of the array represents a parasite, and contains the strain value $x_i$ determining its trait-dependant functions. The system is initialised assigning a starting total number of $R$ and $P$, as well as the value of each element $P[i]$. In our code, the elements $P[i]$ are initialised using a uniform distribution on support $[0, 1]$. Any other distribution can be used to initialise the infected population. Alternatively, they can also be all initialised with the same value. This will only affect the transient dynamics, but not the asymptotic behaviour.

The agent-based dynamics is composed of the following events: *Mutation*, *Exploitation*, *Treatment*, *Mortality*, *Renewal*. Each event is represented by a stochastic process $e$ occurring at a rate $r_e$. During a time-step, each event is simulated once. For each event, a uniformly distributed random number $r_1$ is sampled and compared to $r_i \Delta\tau$, where $\Delta\tau$ is the time-step of the numerical simulation. If $r_1 < r_e \Delta\tau$, then the event $P$ occurs, and the state of the population is updated.

Provided a small enough $\Delta\tau$ (we choose $\Delta\tau = 10^{-5}$), the discrete-time dynamics is expected to converge to the deterministic continuous-time one. In the following, the rates of all the events are presented.

**Mutation.** All parasites have the same probability to mutate. A mutation occurs if $r_1 < \mu_0 \, P \, \Delta\tau$, where $\mu_0$ is the per capita rate of mutation per unit of time. Upon occurrence, a parasite is randomly extracted, and a random variable $\zeta$ is added to its trait value $x$. The random variable $\zeta$ is extracted from a probability distribution $\mathcal{P}(0, \sigma^2)$, with zero mean and variance $\sigma^2$. If the mutated trait value would cross the borders $[0, 1]$ of the trait space, the mutation is rejected and the individual retains its original trait value, that is reflecting boundary conditions are considered.

The assumption of *zero* mean implies that mutations are unbiased, that is the trait space is explored with same probability in any direction. The code provided employs a uniform probability distribution, but the results are expected to hold for any common choice of distribution. For small values of $\sigma^2$, the accumulation of such mutations leads the individuals to perform a random walk exploration of the adaptive space (2). In the deterministic limit, the above process is captured by the local diffusion operator $\nabla^2$, with diffusive coefficient $\mu = \frac{\sigma^2}{2}\mu_0$, and no-flux boundary conditions, as reported in the main text.

**Exploitation.** Each parasite $P$ has a probability of infection that depends on its own trait value $x_i$. In order to simulate this event, we first sample at random a parasite. Then, exploitation occurs if $r_1 < \beta(x_i)\, P\, R\Delta\tau$, where the trait-dependent exploitation rate $\beta(x_i)$ is computed according to Eq. 5 of the main text. Upon occurrence of the event, resource $R$ is decreased, and a new parasite is appended to the array $P$, with trait value equal to $x_i$.

**Mortality.** Resource and parasites are removed proportionally their mortality rate $\delta^{R,P}$. Similarly to the exploitation event, the randomly sampled parasite $P[i]$ will die if $r_1 < \delta_0^P + \delta_1^P(x_i)\ \Delta\tau$. Instead, a resource unit will die if $r_1 < \delta^R\, R\, \Delta\tau$.

**Treatment.** This event is analogous to the previous one, upon replacing $\delta_1^P(x_i)$ with $\gamma\, \phi(x_i)$, where $\gamma$ is the treatment application rate, and $\phi(x_i)$ is the trait-dependent efficacy, computed according to Eq. 6 of the main text. Then, the total number $R$ is increased by one if $\zeta = 1$.

**Renewal.** Resource is renewed at constant rate. Hence, if $r_1 < \theta\Delta\tau$, then the total number $R$ is increased by one.

### 4 Relaxing the linear assumptions

The linear assumption on the treatment efficacy $\phi(x)$ and on the Exploitation-Mortality trade-off $\beta(x)$ presented in the text have been introduced in order to allow analytical calculations of the optimal type and of the application rates thresholds. However, in a more realistic set-up, those relationships are likely to be non-linear and saturating (3). Here we provide a comparison to the main output of our model (Fig. 4 of the main text), obtained when either the above assumptions are relaxed, thus demonstrating the robustness of the qualitative features of the main result. In the non-linear case, a semi-analytical solution of the problem can be obtained by means of a spectral method (4).

**Non-linear Exploitation-Mortality trade-off**. A common choice for $\beta(x)$ is provided by a saturating, Michaelis-Menten-like function (5; 6). Consistently, here we choose:

$$\beta(x) = \beta_0 + \frac{\beta_1}{C_{\beta_2}} \frac{x}{x + \beta_2}, \tag{31}$$

with $\beta_2$ shape parameter, and $C_{\beta_2} = 2\left(1 + \log \frac{\beta_2}{1+\beta_2}\right)$ normalisation factor.

In the left panel of Fig. 1, we compare the function Eq. (31) for different values of the shape parameter $\beta_2$. The shape parameter $\beta_2$ determines the degree of linearity of the function: the limit $\beta_2 \to 0$ corresponds to an almost flat curve, while for $\beta_2 \to +\infty$ the linear case is recovered. The factor $C_{\beta_2}$ ensures that all curves are normalised to $\beta_0 + \frac{\beta_1}{2}, \forall \beta_2$.

In the right panel of Fig. 1, we plot the relationship between equilibrium resource and application rate, for the linear treatment types employed in Fig. 4 of the main text, and $\beta_2 = 1$. Consistently with the results provided by the linear case, we observe again two main features: the optimal treatment type depends on the application rate; there exist an application rate threshold value, above which the gain in resource changes sensibly. Particularly, we remark that non-linearities trigger a smoother saturation than the linear case.

**Non-linear treatment efficacy.** A general spectrum of non-linear efficacies displaying opposite correlation with the trait variable $x$, can be obtained by the following two equations:

$$\phi(x)^{+} = 1 + \phi_1 \left[ x^{\alpha} - \frac{1}{\alpha + 1} \right] \tag{32a}$$

$$\phi(x)^{-} = 1 + \phi_1 \left[ (1 - x)^{\alpha} - \frac{1}{\alpha + 1} \right]. \tag{32b}$$

with $0 \leq \alpha \leq 1$ and $0 \leq \phi_1 \leq (\alpha + 1)$ shape parameters.

Parameter $\alpha$ determines the degree of linearity of the function, with $\alpha = 1$ corresponding to the linear case presented in the main text, and $\alpha \to 0$ representing highly non-linear cases. Parameter $\phi_1$ determines the degree of correlation with respect to the trait variable $x$ (larger values of $\phi_1$ imply larger correlation). The function $\phi(x)^{+}$ is monotonically increasing, therefore modelling treatments that are positively correlated with the trait variable; the function $\phi(x)^{-}$ is monotonically decreasing, therefore modelling treatments that are negatively correlated with the trait variable.

In the left panel of Fig. 2, we plot the functions for $\alpha = \frac{1}{2}$, and different values of $\phi_1$ (explained in the caption). In the right panel of Fig. 2, we explore the relationship between the equilibrium resource and application rate, for the treatment types corresponding to the left panel. The other parameters are those employed in Fig. 4 of the main text.

Again, we observe that the results are consistent with those provided by the linear case. We therefore expect these relevant features to be qualitative inherent to the epi-evolutionary nature of the system, and to be only quantitatively affected by the specific choice of the treatment efficacy and of the Exploitation-Mortality functions.

### References

[1] M. Abramowitz and I. Stegun, *Handbook of mathematical functions: with formulas, graphs, and mathematical tables*, vol. 55. Courier Corporation, 1964.

[2] J. Y. Wakano, T. Funaki, and S. Yokoyama, "Derivation of replicator–mutator equations from a model in population genetics," *Japan Journal of Industrial and Applied Mathematics*, vol. 34, no. 2, pp. 473–488, 2017.

[3] S. Alizon, "Treating symptomatic infections and the co-evolution of virulence and drug resistance," *bioRxiv*, 2020.

[4] L. Miele, R. Evans, and S. Azaele, "Redundancy-selection trade-off in phenotype-structured populations," *Journal of Theoretical Biology*, vol. 531, p. 110884, 2021.

[5] L. Bolzoni and G. A. De Leo, "Unexpected consequences of culling on the eradication of wildlife diseases: the role of virulence evolution," *The American Naturalist*, vol. 181, no. 3, pp. 301–313, 2013.

[6] S. Alizon, A. Hurford, N. Mideo, and M. Van Baalen, "Virulence evolution

and the trade-off hypothesis: history, current state of affairs and the future," *Journal of evolutionary biology*, vol. 22, no. 2, pp. 245–259, 2009.

### 5 Supplementary Figures

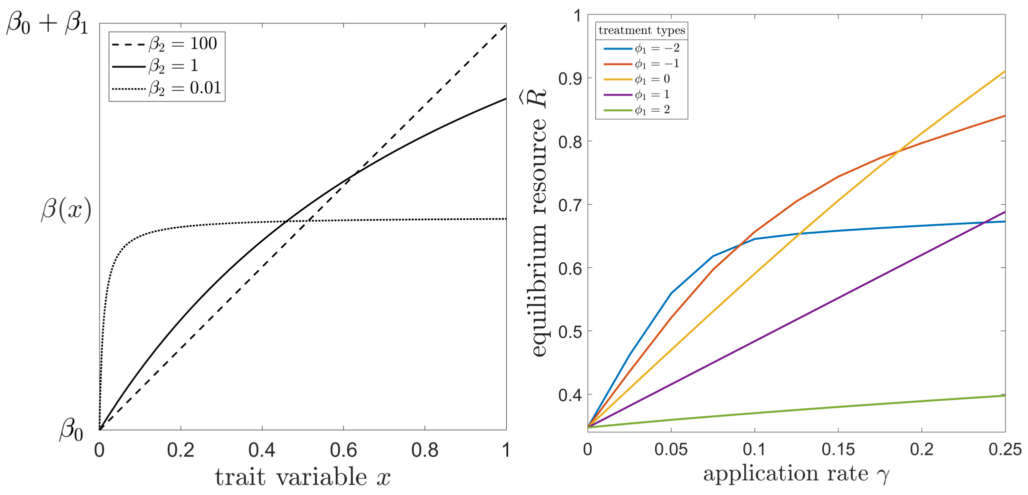


Figure 1: **Non-linear Exploitation-Mortality trade-off.** Left panel: plots of non-linear Exploitation-Mortality curves Eq. (31), for different values of shape parameter $\beta_2$. Right panel: the resource at equilibrium $\widehat{R}$ obtained using five different treatment types, under a non-linear trade-off between exploitation and mortality rates, with $\beta_2 = 1$. Other parameters equivalent to left panel of Fig. 4 of the main text.

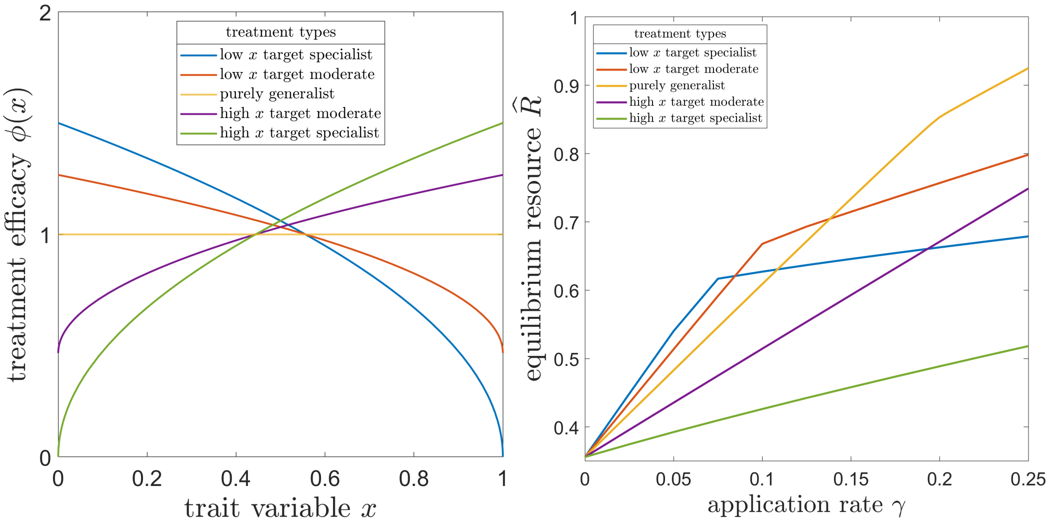


Figure 2: **Non-linear treatment efficacy.** Let panel: examples of non-linear treatment efficacy functions with $\alpha = \frac{1}{2}$ and different values of correlation. Blue line: negative correlation with $\phi_1 = 1.5$, that is a specialist treatment focusing on low $x$ trait values; Red line: negative correlation with $\phi_1 = 0.8$, that is a moderate treatment focusing on low $x$ trait values; Yellow line: purely generalist treatment with $\phi_1 = 0$; Purple line: positive correlation with $\phi_1 = 0.8$, that is a moderate treatment focusing on high $x$ trait values; Green line: positive correlation with $\phi_1 = 1.5$, that is a specialist treatment focusing on high $x$ trait values. Right panel: the resource at equilibrium $\widehat{R}$ obtained using the five treatment types displayed in the left panel, as function of application rate $c$ (the y-axis is normalised with respect to the parasite-free resource $\widehat{R}_0$). Other parameters equivalent to left panel of Fig. 4 of the main text.
